# The essential role of feedback processing for figure-ground perception in mice

**DOI:** 10.1101/456459

**Authors:** Lisa Kirchberger, Sreedeep Mukherjee, Ulf H. Schnabel, Enny H. van Beest, Areg Barsegyan, Christiaan N. Levelt, J. Alexander Heimel, Jeannette A. M. Lorteije, Chris van der Togt, Matthew W. Self, Pieter R. Roelfsema

## Abstract

The segregation of figures from the background is an important step in visual perception. In primary visual cortex, figures evoke stronger activity than backgrounds during a delayed phase of the neuronal responses, but it is unknown how this figure-ground modulation (FGM) arises and whether it is necessary for perception. Here we show, using optogenetic silencing in mice, that the delayed V1 response phase is necessary for figure-ground segregation. Neurons in higher visual areas also exhibit FGM and optogenetic silencing of higher areas reduced FGM in V1. In V1, figures elicited higher activity of vasoactive intestinal peptide-expressing (VIP) interneurons than the background, whereas figures suppressed somatostatin-positive interneurons, resulting in an increased activation of pyramidal cells. Optogenetic silencing of VIP neurons reduced FGM in V1, indicating that disinhibitory circuits contribute to FGM. Our results provide new insight in how lower and higher areas of the visual cortex interact to shape visual perception.

## Introduction

Neurons at early processing levels of the visual system initially analyze the visual scene in a fragmented manner. They carry out a local analysis of the image elements in their small receptive fields (RFs) and feed the visual information forward to higher visual areas (HVAs). Neurons in HVAs have larger RFs and integrate information to represent increasingly abstract features of the visual scene, including object category and identity (Desimone and Duncan, 1995; Yamins and DiCarlo, 2016). However, there are many images for which the analysis is not complete when information has reached the HVAs (Kar et al., 2019). It has been hypothesized that these images require the recirculation of activity back to lower areas through recurrent connections, which include feedback connections from higher to lower visual areas and horizontal connections between neurons within the same cortical area. These recurrent routes are associated with additional synaptic and conduction delays, so that the recurrent influences usually are not expressed when visual cortical neurons are initially activated by the stimulus, but during a later phase of their response (Lamme and Roelfsema, 2000).

One of the proposed roles for recurrent processing is that it supports perceptual organization, the process that groups image elements of behaviorally relevant objects and segregates them from other objects and the background (Roelfsema, 2006). An advantage of recurrent processing for perceptual organization is that neurons in low-level visual cortical areas have small RFs so that they represent the segmentation results at a high spatial resolution. Figure 1A illustrates a few example figure-ground images that induce perceptual organization; a subset of the image elements forms a figure, which is segregated from the background. Studies in monkeys demonstrated that neuronal responses in the primary visual cortex (area V1) that are elicited by figures are stronger than those elicited by background regions, even if the image elements in the RF are the same (compare the RF for stimulus 1 vs. 2 and 3 vs. 4 in Figure 1B) (Chen et al., 2014; Lamme, 1995; Lamme and Roelfsema, 2000; Poort et al., 2016; Self et al., 2019). This relative enhancement of neural activity on figures is known as figure-ground modulation (FGM). FGM has been measured with fMRI and EEG in the visual cortex of human participants (Scholte et al., 2008) and a study in a human subject documented that image elements of a figure also elicit stronger spiking activity than background elements in low-level areas of the visual cortex (Self et al., 2016).

**Figure 1.**
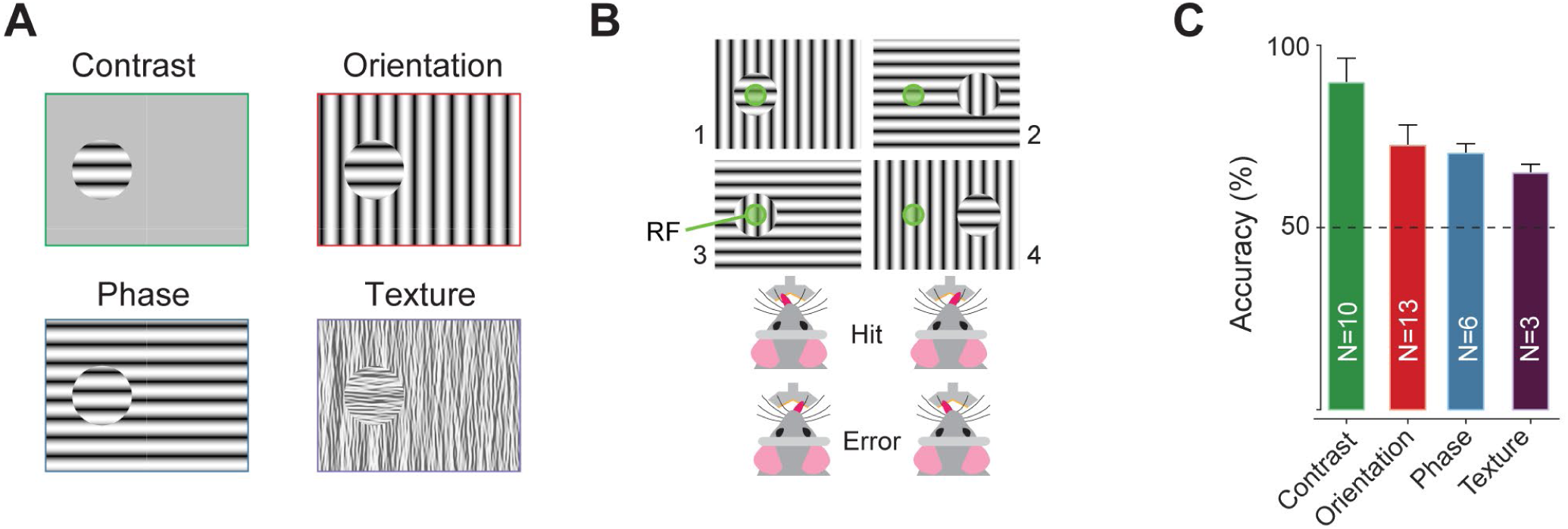
Mice segregate different types of figures from a background. (A) Stimuli that induce figure-ground perception. The stimulus either contained a grating that differed from the background in orientation (top right) or phase (bottom left). For comparison, we presented a circular grating on a grey background (“contrast”, top left) that did not require figure-ground segregation. We also presented a texture stimulus with an orientation difference between figure and ground (bottom right). (B) The mice reported the figure-location by licking a spout on the corresponding side. The image elements falling in the RF (green circle) were identical in the figure and background conditions so that differences in neuronal activity could only be caused by the context determined by stimulus regions outside the RF. (C) Average accuracy of the mice when performing the task with different stimuli. Error-bars, standard deviation across mice. The number of mice per experiment is indicated within each bar.

FGM is a contextual effect (it originates from outside the neurons’ RFs) that occurs later than the initial response elicited by stimulus onset, suggesting that it represents a recurrent influence on the neuronal response, which originates from HVAs (Klink et al., 2017; Lamme and Roelfsema, 2000). In accordance with an important role of feedback connections, FGM in V1 is stronger when a monkey pays attention to a figure (Poort et al., 2012) or if it enters into the subject’s awareness (Supèr et al., 2001). However, direct evidence for a causal role of feedback connections in FGM has been lacking. Specifically, it is unknown whether FGM is just an epiphenomenon of processing in HVAs that feed back to V1, or if the delayed V1 response phase is necessary for perception. Hence, three of our aims are to test whether the late V1 response is necessary for figure-ground perception, whether FGM occurs in HVAs and if HVAs are the source of FGM in V1.

A fourth aim was to gain insight into the roles of interneurons in figure-ground segregation. A previous study proposed that the activity of somatostatin-expressing inhibitory neurons (SOM-neurons) is a candidate mechanism for background suppression (Adesnik et al., 2012). SOM-neurons inhibit the dendrites of pyramidal neurons and respond strongly to homogeneous image regions that are likely to be part of the background. Another study (Zhang et al., 2014) suggested a mechanism by which feedback connections could increase the activity for relevant representations in lower areas via vasoactive intestinal peptide-positive neurons (VIP-neurons). VIP-neurons are interneurons that inhibit SOM-cells, and they thereby disinhibit the cortical column (Harris and Shepherd, 2015; Karnani et al., 2016; Pi et al., 2013). However, the activity of interneurons elicited by figure-ground stimuli and their involvement in generating FGM in V1 remains to be demonstrated.

Previous studies in freely moving mice demonstrated that mice can perceive figures that differ from the background in orientation or phase (upper right and lower left panels in Figure 1A) (Li et al., 2018; Schnabel et al., 2018). To address our questions, we developed a figure-ground paradigm for head-fixed mice, to have sufficient control over the stimulus position relative to the mouse and simultaneously measure brain activity. Furthermore, we tested the generality of figure-ground perception by also including stimuli where the figure differed from the background in texture (lower right panel in Figure 1A), similar to those used in previous monkey studies (Lamme, 1995).

We report that HVAs are crucial for the generation of FGM in V1 and that figure-ground perception fails if the late V1 response phase is inhibited. Furthermore, our results demonstrate that figures enhance VIP-cell activity and suppress the activity of SOM-cells. We report that VIP-neurons cause disinhibition, which is more pronounced at the figure location so that optogenetic suppression of VIP-cell activity reduces FGM. These new insights into the neuronal mechanisms for figure-ground perception are likely to generalize to other cases in which the neuronal representations of simple features in lower brain areas and more abstract features in higher brain areas need to interact with each other during perception.

## Results

### Mice can segregate figures from the background using various cues

We trained head-restrained mice to report whether a circular grating (0.075 cycles/degree) or texture pattern was positioned on the left or right side of a display by licking one of two spouts (Figures 1A,B). We rewarded correct choices with a drop of liquid reward. In the initial training phase, mice learned to report the location of figures on a grey background. We then gradually increased the contrast of the background grating until it equaled the contrast of the figure grating and the only remaining difference was orientation or phase. Mice reached an accuracy of 90±7% (mean±s.d.; N=10 mice) in the task with a grey background, 72±6% (N=13 mice) if the figure differed in orientation from the background, and 70±3% if it differed in phase (N=6 mice) (Figure 1C). The accuracy of the mice was significantly above chance level (ps<0.001 for all mice and tasks, Bonferroni corrected one-sided binomial test; Table S1). To test generalization, we presented a new figure-ground stimulus with orthogonally oriented textures (Figure 1A, bottom right) and observed immediate generalization with an average accuracy of 65±2% (N=3 mice, ps<0.001 for all mice). The demonstration that head-restrained mice can report figure-ground structure elicited by various cues implies that the many powerful methods available to measure and change neuronal activity in this species can be used to study the neuronal mechanisms of perceptual organization.

### Figure-ground modulation in V1 is evoked by various visual cues

To examine V1 activity elicited by these different figure-ground stimuli, we recorded multi-unit activity (MUA) with multichannel silicon probes (N=198 recording sites in 13 electrode penetrations in 6 awake mice, passively viewing the stimuli). For the figure-ground stimuli, we ensured that the stimulus in the RF was identical in the figure and ground conditions (Figures 1B, 2A), so that activity differences could only be caused by contextual information from stimulus regions outside the classical RF. For comparison, we also presented contrast stimuli (Figure 1A, upper left) where the RF either fell on a grating or the grey background. We quantified FGM for individual recording sites with d-prime, a measure of reliability of the difference between the figure and ground response on single trials (time-window 100-500ms after stimulus onset). Activity elicited by orientation, phase and texture-defined figures was stronger than that elicited by backgrounds, despite the sensory stimulus in the RF being identical (Figure 2B; Figure S1 shows responses of example recording sites). We used linear mixed-effects models (Methods) to account for the fact that recording-sites from the same penetration typically have higher correlations than those from different penetrations (Scheffe, 1956) and observed significant FGM for the three types of figure-ground stimuli (all ps<0.001). The average d-prime was 1.65 for the contrast stimulus (indexing the reliability of the visual response) and it was 0.44, 0.22 and 0.24 for the orientation-defined, phase-defined and texture stimuli, respectively (indexing FGM). FGM was stronger for orientation-defined figures than for figures with a phase or texture difference (ps<0.001, Bonferroni corrected post-hoc t-tests). To determine FGM-latency, we subtracted V1 activity elicited by the background from that elicited by the figure and fitted a curve to the response difference (lower panels in Figure 2B; see Methods). We measured a visual response latency of 43ms with the contrast-defined stimulus. FGM latencies for all figure-ground stimuli were delayed relative to the visual response latency and only occurred during a later processing phase (all ps<0.001, bootstrap test). The latency of FGM for orientation defined figure-ground stimuli was 75ms. Latencies for the phase-defined (91ms) and texture figure-ground stimuli (92ms) were, in turn, longer than for the orientation-defined figure-ground stimulus (both ps≤0.001, bootstrap test). We also measured FGM in V1 of four mice performing the figure detection task (Figure S2). The strength of FGM on correct trials was similar to that during passive viewing and even to that on error trials.

**Figure 2.**
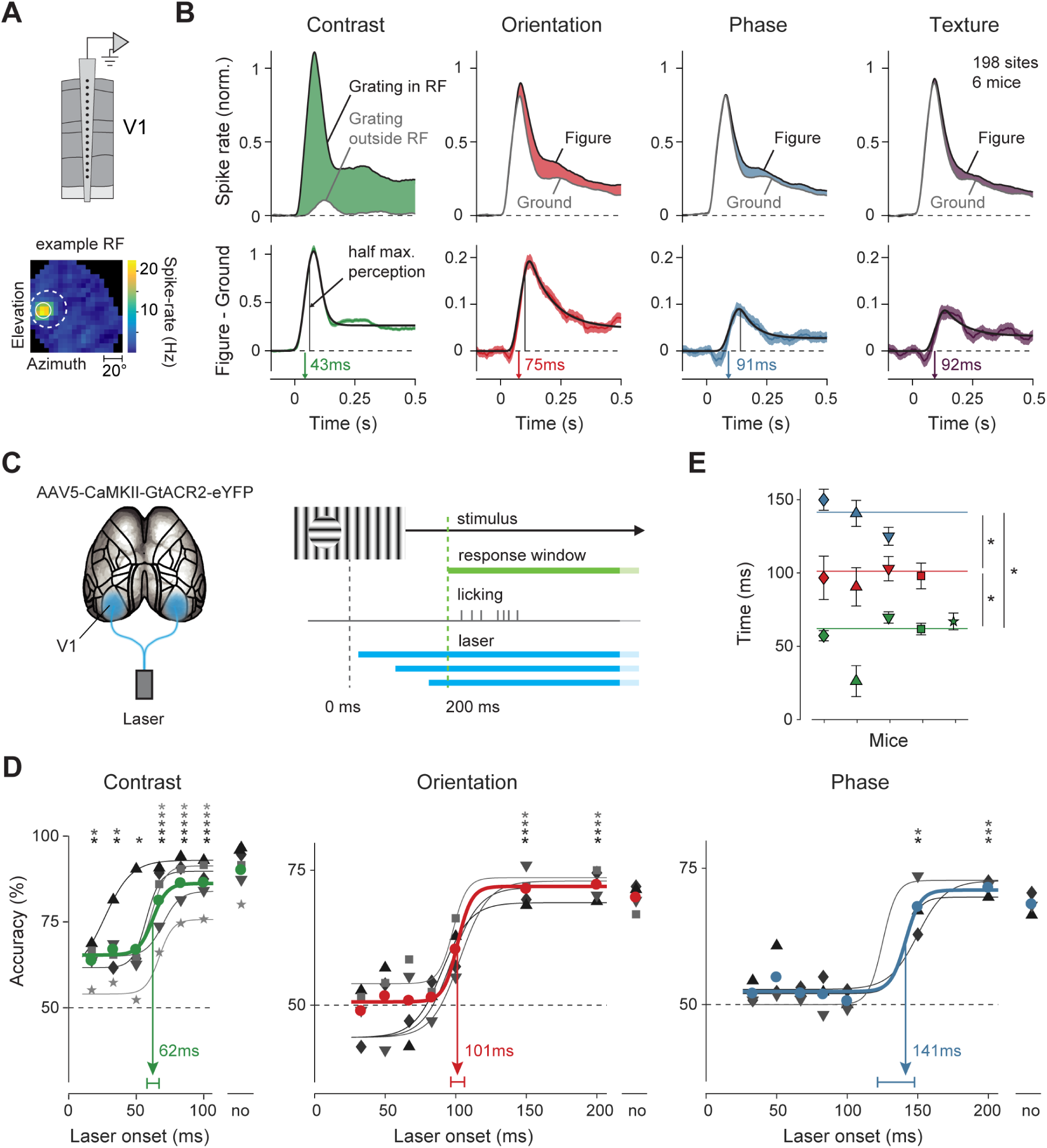
Figure-ground perception relies on late V1 activity. (A) Silicon probe recordings in V1 and an example RF determined with a mapping stimulus (Methods). White continuous circle, estimated RF-boundary. Dashed circle, figure location. (B) Average MUA-response elicited by a contrast-defined figure and the figure-ground stimuli (198 recording sites in 6 mice). The lower panels show the difference in activity elicited by figure and ground (FGM) (visually driven response for the contrast stimulus) and a curve that was fitted to determine FGM latency (visual response latency for the contrast stimulus; see Methods). The black vertical lines illustrate the V1-processing time required for half-maximal accuracy (from panels D,E). (C) We optogenetically inhibited V1 activity in both hemispheres at different delays after stimulus appearance using blue laser light. The mice reported figure location after 200ms by licking. (D) Accuracy of five mice in the tasks, as function of the onset time of optogenetic V1 silencing. The data points on the right of the graphs are the accuracies on trials without silencing. The data of individual mice are shown as grey/black symbols and curves are fits of logistic functions. Colored curves, fits to the average accuracy across mice. The inflection point of the curve was taken as measure of latency (the 95% confidence interval is shown above the abscissa). Asterisks above the plot indicate when the accuracy of the individual mice was above chance level (p<0.05, Bonferroni-corrected binomial test). (E) V1 processing time necessary to reach half-maximal performance in the contrast detection task (green) and for orientation-defined (red) and phase-defined (blue) figure-ground stimuli. Symbols show data from individual mice (same symbols per mouse as in D) and the horizontal lines show the average latencies. Error bars, s.e.m. determined by bootstrapping.

### Late V1 activity is necessary for figure-ground segregation

FGM in V1 occurs during the late, sustained processing phase. To determine if late V1 activity is necessary for figure-ground perception, we inhibited V1 neurons at different latencies while mice were performing the figure-ground segregation task. We expressed the inhibitory opsin GtACR2, a light activated chloride channel (Govorunova et al., 2015), bilaterally in V1 pyramidal neurons in five mice. Blue light quickly and reliably inhibited the neuronal activity across all cortical layers (Figures S3F-H). We varied the onset of the blue light across trials so that we could examine the phases of the V1 response that are essential for reliable performance (Resulaj et al., 2018) (Figure 2C). In the contrast detection task, silencing from 33ms after stimulus onset onwards, which abolished the entire V1 response, reduced the accuracy of the mice (Figure 2D) (Bonferroni corrected 1-sided binomial test, p<0.05 in each mouse). In most animals, accuracy remained above chance level, which suggests that a relatively low accuracy in the contrast detection task can be maintained by brain regions outside V1, although we cannot exclude the possibility that V1 silencing was incomplete in some of the mice. The accuracy quickly recovered when we postponed optogenetic silencing until after the visually driven V1 response, reaching its half maximal value after 62ms (95%-confidence interval 58-67ms) (Figures 2D,E). In the orientation-defined figure-ground task, silencing of V1 reduced the accuracy to chance level and it only recovered to its half-maximal value when V1 silencing was postponed to 101ms (95%-confidence interval 97-106ms). The required processing time was even longer (141ms) if the figure was defined by a phase-difference (95%-confidence interval 121-148ms; ps<0.05 for latency differences between all conditions, Figure 2E). We also measured the minimal reaction time (mRT) for the different stimuli, as the 10ms bin of the reaction time distribution at which the mice made more correct than erroneous licks (Kirchner and Thorpe, 2006). For contrast-defined figures, the mRT was 245ms and it was 255ms and 315ms for the orientation- and phase defined figure-ground stimuli, respectively (Figures S3A-C). We found that mRTs for phase-defined figures were longer than for orientation-defined figures (paired t-test, p<0.05; Figure S3B), in accordance with the longer minimally necessary V1 processing time for these stimuli.

We ensured that the blue light itself did not cause interference by carefully shielding the mouse’s eyes from the blue light that shone on the cortex and placing a second blue LED below the mouse’s head that flashed at random intervals, which the animal learned to ignore (see Methods). Furthermore, shining blue light on cortical areas not expressing the optogenetic channel did not reduce the accuracy in the task (Figures S3D,E). We conclude that the initial V1 response transient suffices for accurate performance in the contrast detection task, whereas the late V1 response phase is necessary for figure-ground segregation, supporting the hypothesis that it is read out by brain structures that select the appropriate motor response (Roelfsema and de Lange, 2016). The latencies derived from the optogenetic experiment were systematically longer than the neuronal latencies (compare neuronal FGM latencies to time of half max. accuracy; black lines in the lower panel of Figure 2B), suggesting that downstream areas need to integrate V1-FGM during a longer epoch to support accurate performance.

### Figure-ground modulation is present in multiple higher cortical areas

The late onset of FGM in V1 suggests that it may be a result of recurrent interactions between V1 and HVAs. To examine neuronal activity across the cortical areas that could provide feedback to V1 (Gamanut et al., 2018; Oh et al., 2014; Osakada et al., 2011; Wang et al., 2011), we used wide-field imaging in nine Thy1-GCaMP6f mice with a transparent skull (Dana et al., 2014; Guo et al., 2014). We first measured the retinotopy to determine the boundaries between the areas and to align them with the Allen Brain Map (see Methods, Figures S4, S5D) and investigated FGM elicited by orientation-(N=9 mice) and phase-defined (N=5 mice) figure-ground displays, during passive viewing. We used the measured retinotopy to transform the V1 activity profile into visual field coordinates and we observed that the entire visual field region occupied by the figure was labeled with enhanced activity, with a protracted time-course caused by the dynamics of the calcium-sensor (time-window 150-300ms; paired t-test, p<0.001) (Figures 3A,B, S5A,B). Orientation- and phase-defined figures also elicited more activity than the background in areas LM, AL, RL, A, AM, PM and retrosplenial cortex (RSP; Figure 3C) (ANOVA for d-primes across mice, followed by post-hoc t-tests, all ps<0.05). To examine if FGM increases during active perception, we trained five of these mice to perform the orientation figure-ground task during wide-field imaging. FGM during correct figure-detection trials was stronger than during passive viewing in areas V1, AL, A, AM, PM and RL (two-way ANOVA with factors area and task; interaction, F_9,36_ = 10.6, p<0.001; post-hoc tests for V1, AL, A, AM, PM and RL, ps<0.05; Figure 3D), in accordance with a previous study demonstrating that activity in some of these areas increases when mice engage in a visual task (Pho et al., 2018). We also compared the activity between correct and erroneous trials. Just as in our electrophysiological experiments, FGM was present when mice made an error. In some visual areas there was a trend for stronger FGM for trials with correct responses (area A, p=0.06, two-sided t-test, Figure S5C) but this effect was not significant. These results therefore indicate that FGM is a robust phenomenon that occurs in different behavioral states and even during erroneous trials.

**Figure 3.**
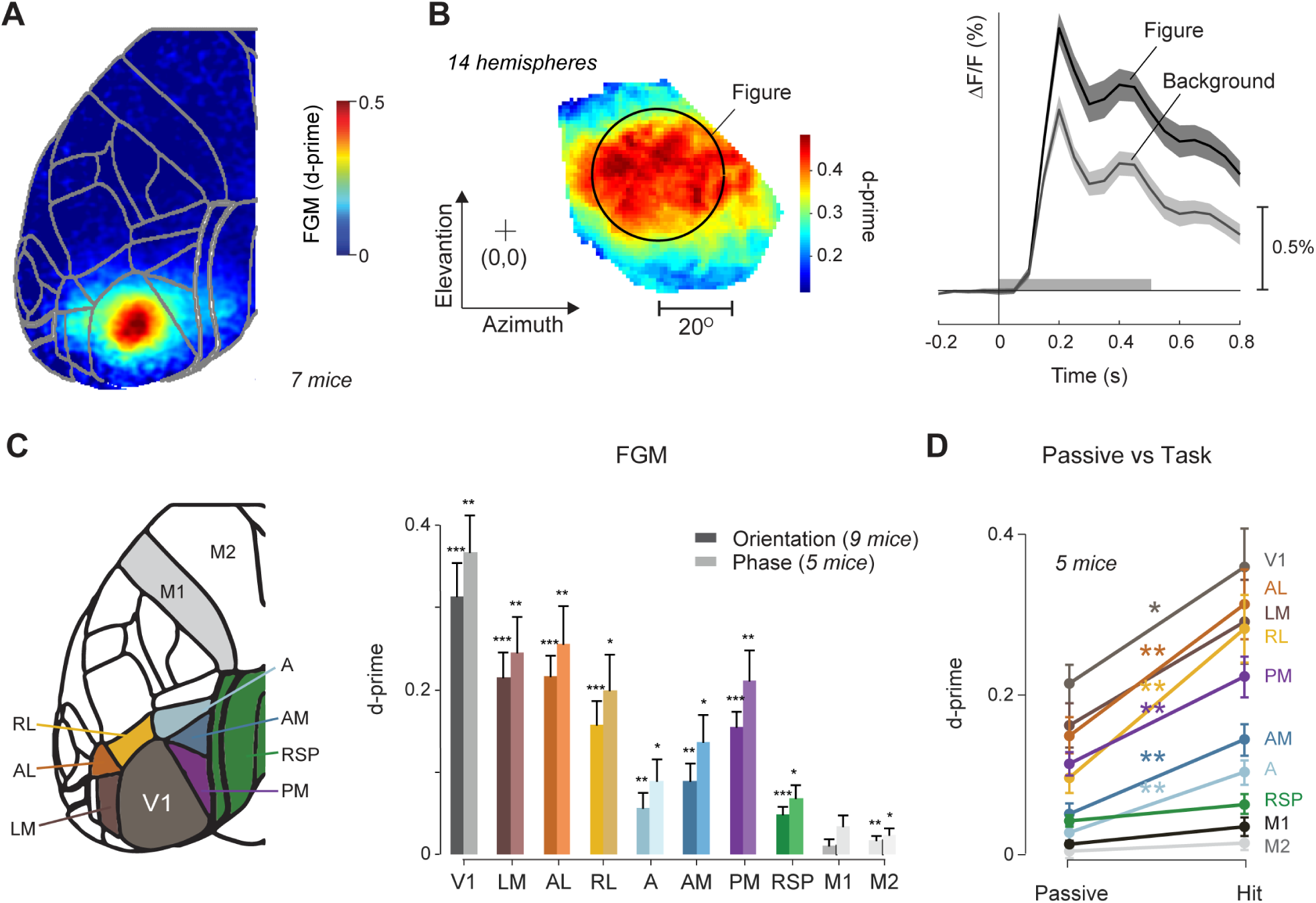
FGM across areas of the visual cortex measured with wide-field imaging. (A) Strength of FGM for orientation-defined figures quantified using the d-prime during passive viewing, averaged across all hemispheres at 150-300ms after stimulus onset. We used 7 mice, 2 hemispheres per mouse. (2 mice were not used in this analysis because we used a different figure position in these animals). Data of individual mice is presented in Figure S5A. (B) left, the average V1 d-prime values over 14 hemispheres from 7 mice elicited by an orientation defined figure-ground stimulus transformed into visual coordinates, based on the retinotopy. Data from the right visual-field was reflected horizontally and projected into the left visual-field prior to averaging. The black circle illustrates the outline of the figure appearing on the screen in front of the mouse. Plus symbol, center of the visual field. The colored region indicates the extent of the visual field which could be reliably mapped in at least 5 mice per hemisphere. right, Average time-course of the response of V1 pixels falling in the figure representation (black circle in the left panel) when presented with an orientation-defined figure (black curve) or background (grey curve) (for the response phase-defined stimuli, see Figure S5B). The shaded area denotes s.e.m. across hemispheres. (C) left, We aligned the brains of individual mice to the Allen brain common coordinate framework (Methods). right, D-prime (averaged across all pixels within each area) for orientation and phase defined figure-ground stimuli in passively viewing mice. *, p<0.05; **, p<0.01; ***, p<0.001, post-hoc t-tests. Error bars denote s.e.m. (D) D-prime for orientation defined figure-ground stimuli during passive viewing and hits in the figure-detection task. Error bars denote s.e.m. (N=5 mice). FGM was increased during hits in areas V1, AL, RL, PM, AM and A; *, p<0.05; **, p<0.01.

Wide-field imaging pools calcium signals across multiple cellular compartments, with possible contributions of axons that originate in other brain regions and the method emphasizes layer 1 activity (Allen et al., 2017). We therefore also examined the calcium signals of cell bodies of excitatory neurons within area V1 in five mice using two-photon imaging (Figure 4A). The mice were injected with the cell specific virus AAV1-CaMKII-GCaMP6f-WPRE-SV40 and passively viewed stimuli with figures defined by an orientation difference. Just like in the electrophysiological (Figure 2B) and wide-field recordings (Figure 3), V1 responses elicited by figures were elevated relative to those elicited by the background (Figures 4C-E; time-window 300-1500ms; linear mixed-effects model, see Methods; 1122 cells, p<0.001).

**Figure 4.**
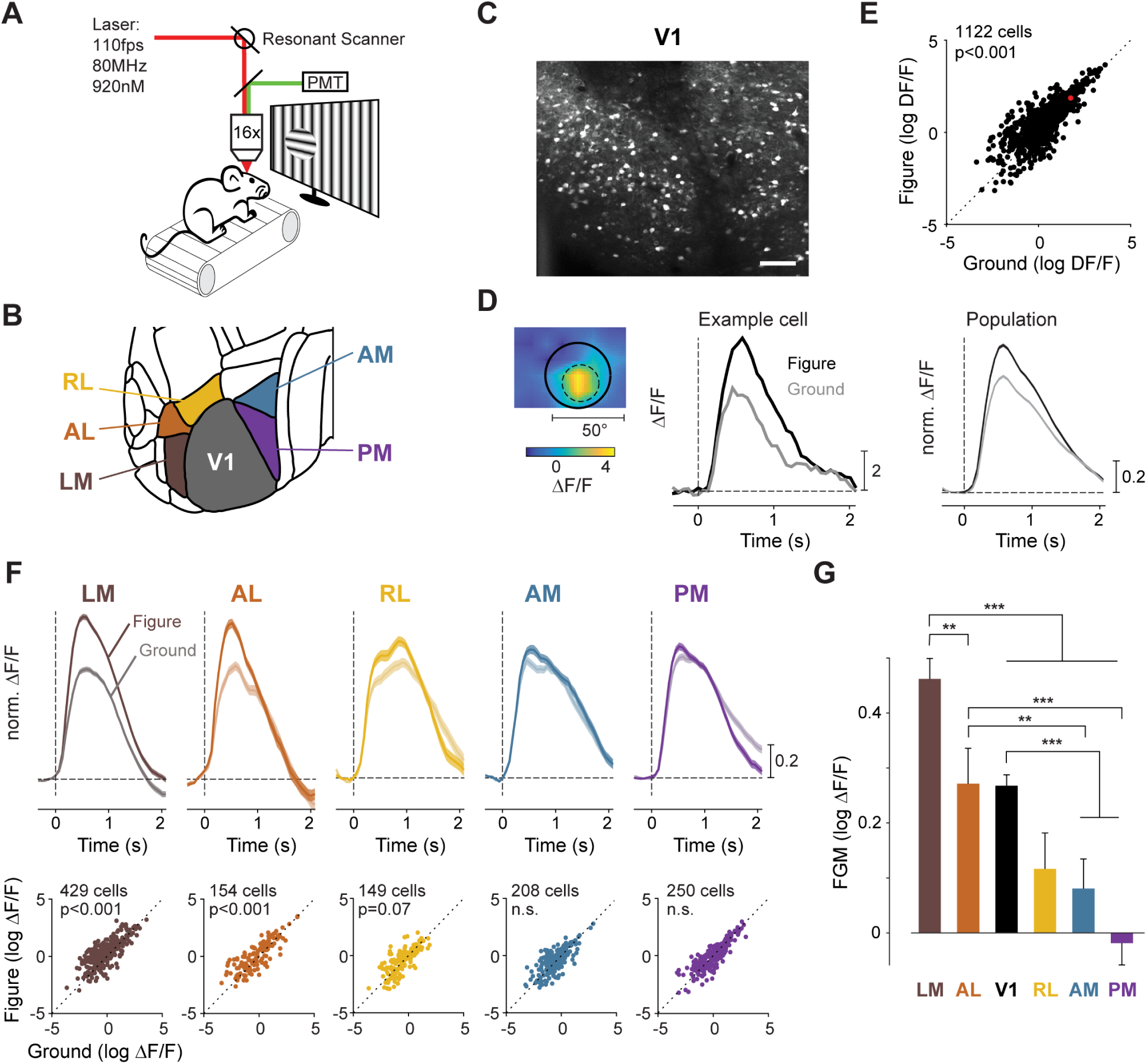
Ca^2+^-response of pyramidal cells in higher visual areas during figure-ground segregation. (A) We used two-photon imaging to measure the Ca^2+^-responses of pyramidal neurons in five Thy1-GCaMP6f mice, additionally injected with AAV1-CaMKII-GCaMP6f-WPRE-SV40 in the visual cortex. The mice were head-fixed and could run on a treadmill while passively viewing orientation-defined figure-ground stimuli. (B) Organization of visual areas in the mouse cortex. (C) We visualized the regions with calcium responses by computing correlation images; luminance values represent the average correlation between a pixel and its 8 neighbors across 3,000 frames. Scale bar, 100µm. (D) We determined the cell-body RF (dashed circle) and placed the figure (continuous circle) or the background in the RF. The middle and right panels show the activity of an example pyramidal neuron and the average activity of the population of 1122 pyramidal cells in V1 elicited by the figure-ground stimuli, after normalization to the maximum of the figure condition. Average activity is shown in normalized units (n.u.). The shaded area shows ± s.e.m but the shading is difficult to see given its small magnitude. (E) The activity of pyramidal cells in V1 in a time window from 300-1500ms after stimulus onset. As the data were positively skewed we plot log (ΔF/F). Figures elicited stronger responses than the background. Red symbol, example cell of panel D; note that the log-transform causes the symbol to be just above the diagonal, in spite of substantial FGM. (F) Ca^2+^-responses of cell bodies of excitatory cells in higher areas LM, AL, RL, AM and PM, in the same mice of panel E. Upper panels, normalized population activity of excitatory cells evoked by figures (saturated colors) and backgrounds (less saturated colors) in normalized units (n.u.). Shaded areas, s.e.m. Lower panels, log (ΔF/F) of individual excitatory cells (time window 300-1500ms after stimulus onset) evoked by figures (y-axis) and the background (x-axis). (G) Comparison of FGM in six visual areas based on log-transformed activity. FGM was strongest in LM, followed by AL and V1, weaker in RL and weakest in AM and PM (Bonferroni corrected post-hoc tests; **, p<0.01; ***, p<0.001).

In the same mice, we recorded calcium signals of cell bodies in the cortical HVAs: LM, AL, RL, AM and PM (Figure 4B). FGM was particularly strong in LM (Figure 4F), and its strength in AL was comparable to that of V1 and it was weaker in RL. These results confirm that neurons in multiple HVAs exhibit FGM (linear mixed-effects models in five mice; LM: 429 cells, F_1,856_=154, p<0.001; AL: 154 cells, F_1,306_=17.9, p<0.001; RL: 149 cells, F_1,296_=3.2, p=0.07). FGM in AM and PM-neurons was even weaker during passive viewing (both ps>0.1; see Figure 4G for comparisons between areas). We note, however, that the wide-field imaging results revealed AM and PM as areas in which FGM increases during task performance (Figure 3D). Furthermore, the comparison of two-photon and widefield imaging results supports the view that widefield imaging emphasizes layer 1 activity (Allen et al., 2017), because LM cell bodies had stronger FGM than V1 cell bodies and they provide strong input into layer 1 of area V1 (Yang et al., 2013).

### Higher visual areas contribute strongly to figure-ground modulation in V1

Figures cause stronger activity than backgrounds in HVAs. We next investigated if feedback from HVAs could be a source of the increased V1 activity that is elicited by figures, during the delayed response phase. Alternative possibilities are that FGM is generated locally, within V1 (Li, 1999) or that it reflects an influence from subcortical visual areas, mediated via the thalamus. To examine the contribution of cortical feedback to FGM in V1, we silenced HVAs (Figure 5A). We injected AAV1-CaMKII-stGtACR2-FusionRed medially (in areas PM, AM and A) and laterally from V1 (LM, RL and AL) in Thy1-GCaMP6f mice (Dana et al., 2014). The virus encodes a soma-targeted version of the inhibitory opsin GtACR2 (Mahn et al., 2018), which was expressed in excitatory neurons. We ensured that the virus did not spread into V1 by identifying the area borders with population RF mapping and targeted the viral injections at a minimal distance of 0.5mm from the V1 border (Figures 5B,C). We used laminar silicon probes to record V1 activity elicited by figure-ground stimuli, and optogenetically inhibited neuronal activity in the HVAs on 50% of the trials, from 200ms before stimulus onset until 100ms after stimulus offset. Optogenetic inhibition did not have a systematic influence on the early peak response of V1 neurons (time-window 0-100ms, Figure S6A), but it reduced the late V1 responses elicited by figures and backgrounds (Figure S6C). This V1 activity reduction was more pronounced for figures than for backgrounds, decreasing FGM by 55%, 100%, and 79% for the orientation defined, phase-defined and textured figure-ground stimuli, respectively (Figures 5D-L, S6D) (5 mice, 8 penetrations, 109 recording sites, time-window 100-500ms, linear mixed-effects model, p<0.001 for orientation, p<0.05 for phase and texture defined figures). Hence, feedback from HVAs enhances the representation of the figure in V1 more than the representation of the background and it thereby accounts for a large fraction of the FGM in V1.

**Figure 5.**
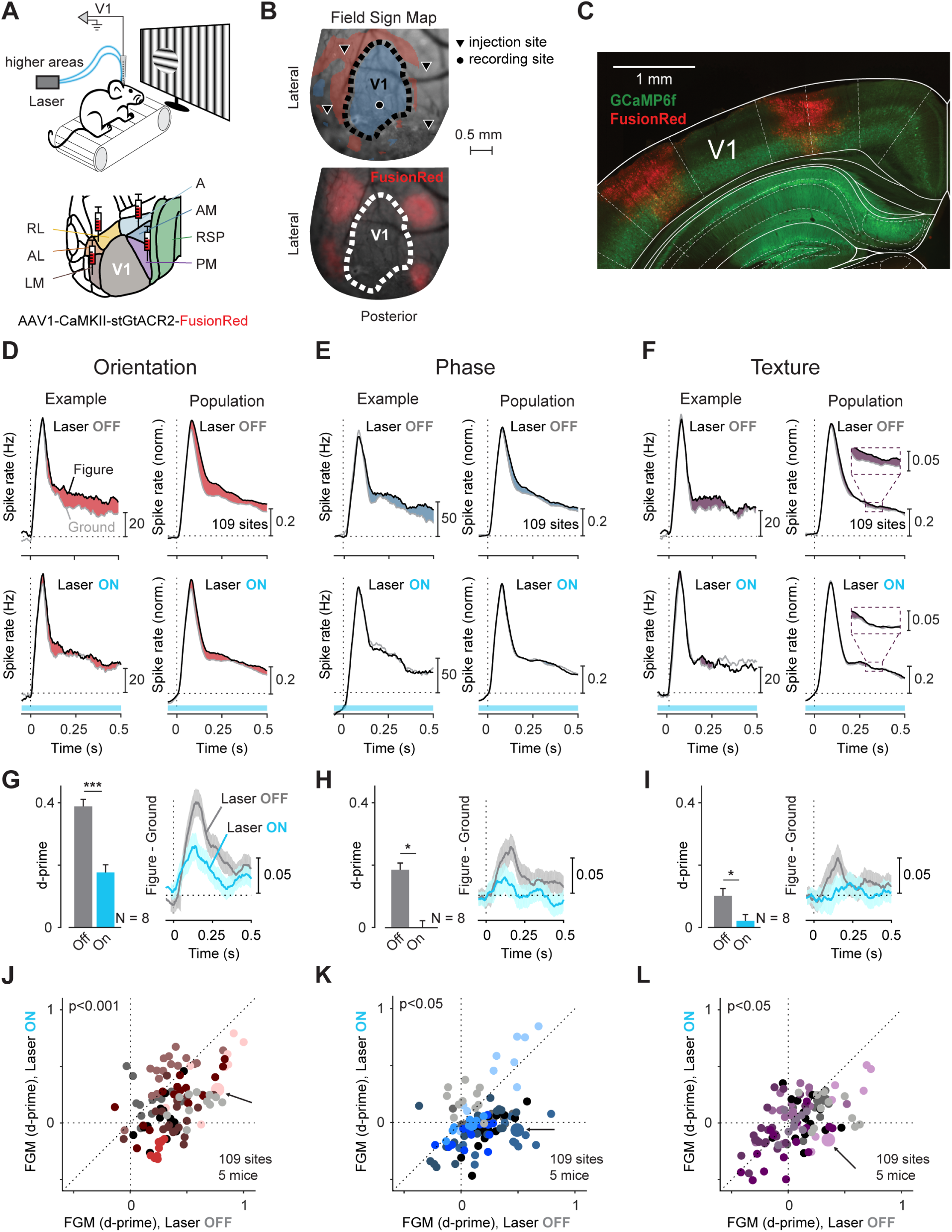
Contribution of feedback from higher visual areas to FGM in V1. (A) Mice were head-fixed in front of a large screen on a treadmill. We injected AAV1-CaMKII-stGtACR2-FusionRed, encoding for the inhibitory chloride channel stGtACR2 in HVAs (four injection sites along a ring around V1) and inhibited activity by shining laser light on HVAs lateral and medial to V1 while recording V1 activity with laminar electrodes. (B) Upper panel, Example field-sign map generated using population RF mapping, overlaid on the visual cortex (Methods). Blue (red) regions have a mirror-inverted (non-inverted) retinotopy. The black circle shows the location of an example electrode penetration, guided by the field-sign map and the blood vessel pattern. Triangles, virus injection sites, which were at least 500µm from the boundary of V1 (dashed line). Lower panel, expression profile (wide-field fluorescence image). (C) Coronal brain slice showing virus (red) and GCaMP6f expression in the Thy-1 mice (green) aligned to the Paxinos and Watson adult mouse atlas. (D) MUA at an example V1 recording site (left) and across a population of 109 sites (8 electrode penetrations, 5 mice) (right) elicited by an orientation-defined figure (black curve) and the background (grey curve) without (top) and with optogenetic silencing (bottom) of HVAs. Data for population responses was normalized to the laser off condition (Methods). The example site is marked in panel J with an arrow and a larger symbol. (E,F), MUA elicited by phase-defined (E) and texture-defined figure ground stimuli (F). (G-I) left, FGM quantified as d-prime (time window 100-500ms after stimulus presentation) in V1 was lower during optogenetic inhibition of HVAs (linear mixed-effects model, see Methods) for orientation-defined (G), phase-defined (H) and textured figure-ground stimuli (I); *, p<0.05; **, p<0.01; ***, p<0.001. right, Average time course of FGM across 109 sites without (grey curve) and with inhibition of activity in HVAs (blue curve). (J-L) FGM d-prime with (y-axis) and without (x-axis) optogenetic inhibition of HVAs for orientation defined (J), phase-defined (K) and textured stimuli (L). Data of different penetrations are shown in distinct colors (linear mixed-effects model, p<0.001 for orientation and p<0.05 for phase and textured figure-ground stimuli).

### Roles of the different interneuron classes in V1 figure-ground modulation

We next asked how the feedback signal from HVAs might influence activity in the cortical microcircuit in V1 to enhance the activity evoked by figures. Recent studies suggested that the different types of interneurons have unique roles in controlling the activity of the cortical column (Harris and Shepherd, 2015; Tremblay et al., 2016). We therefore examined how the three main interneuron subclasses in V1 respond to figure-ground stimuli and compared their activity to that of pyramidal cells. Feedback axons can enhance V1 activity by directly targeting excitatory neurons, and suppress activity by contacting parvalbumin (PV) or somatostatin (SOM) expressing interneurons (D’Souza et al., 2016; Wall et al., 2016; Zhang et al., 2014) (Figure 6A). A previous study demonstrated that SOM-neurons suppress V1 activity elicited by larger, homogeneous image regions (Adesnik et al., 2012) but their response to figure-ground displays has not yet been studied. Finally, feedback connections to V1 could also disinhibit the cortical column by targeting vasoactive intestinal peptide-positive (VIP) interneurons, which inhibit SOM-cells and thereby cause disinhibition of pyramidal cells (Harris and Shepherd, 2015; Karnani et al., 2016; Pfeffer et al., 2013; Pi et al., 2013; Williams and Holtmaat, 2019; Zhang et al., 2014) (Figure 6A). The relative contribution of these excitatory, inhibitory and disinhibitory motifs to perceptual organization is unknown and modeling studies indicate that the interactions between the cell types might be complex, given the large number of influences between the different cell-types (Dipoppa et al., 2018; Garcia Del Molino et al., 2017). Nevertheless, a few predictions can be made. First, VIP-neurons are predicted to be more active for the figure than for the background. Second, their targets, SOM-neurons should respond more weakly to the figure than to the background, so that FGM should be inverted for these cells.

**Figure 6.**
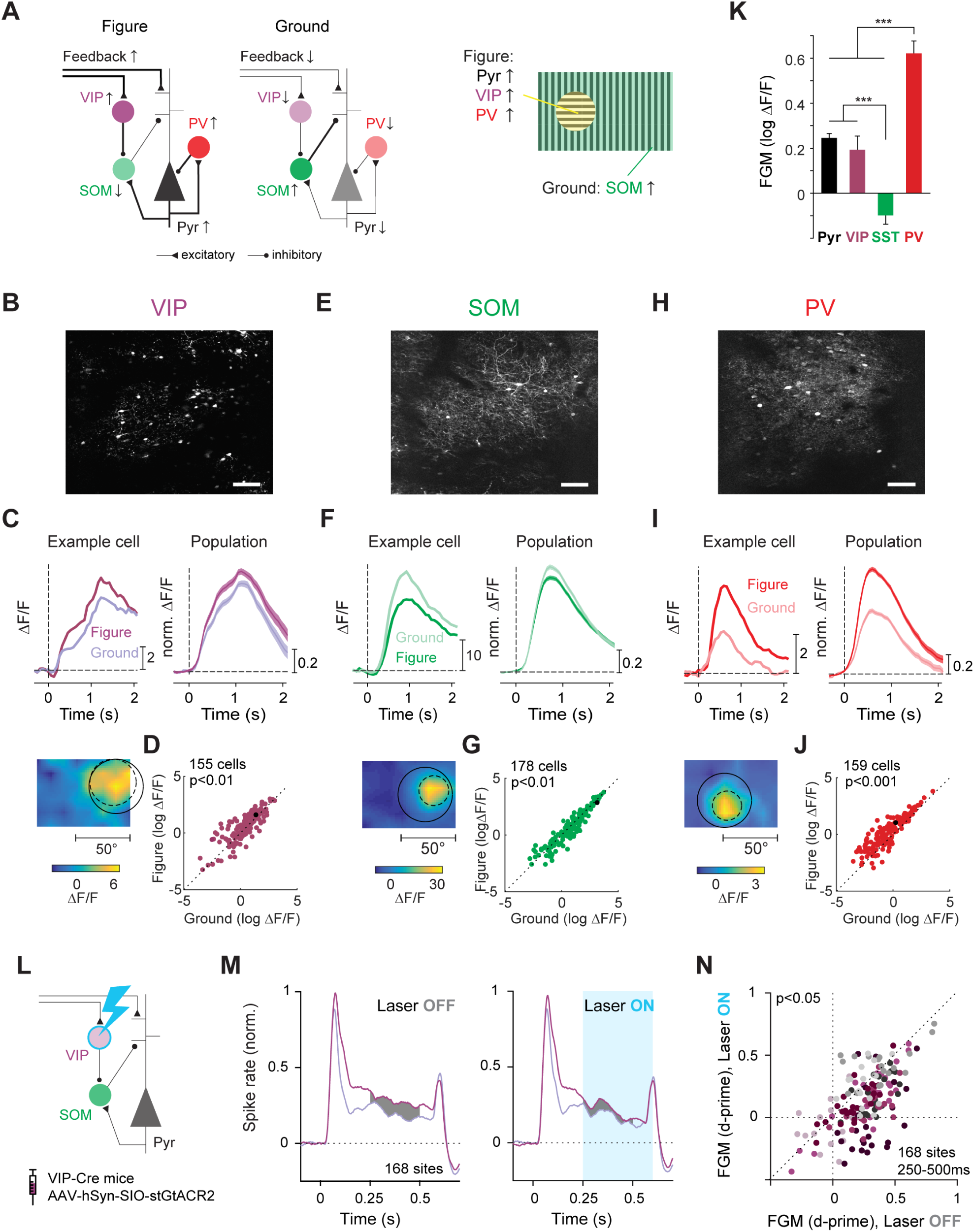
The activity of interneurons in area V1 during figure-ground segregation. left, middle, Previous studies demonstrated that homogeneous image regions cause SOM-neuron mediated inhibition of V1 pyramidal cells (Pyr) (Adesnik et al., 2012), whereas feedback connections target VIP-neurons that inhibit SOM-neurons, thereby disinhibiting the cortical column (Zhang et al., 2014). PV-cells are thought to cause feedforward inhibition and control the gain of the cortical column. right, Our results reveal that figures enhance the activity of pyramidal neurons, VIP-, and PV-cells, whereas SOM-neurons are more active for the background. (B) GCaMP6f expression in an example VIP-Cre mouse induced with AAV9-CAG-Flex-GCaMP6f. (C) Activity of an example VIP-neuron and a population of 155 VIP-neurons in four mice elicited by the figure-ground stimuli. The lower panel illustrates the RF of the example cell. (D) Figures elicited stronger responses in VIP-neurons than the background (linear mixed-effects model, p<0.01). Black symbol, the example cell from panel C. (E) The expression of GCaMP6f in an example SOM-Cre mouse after AAV injection in V1. We only imaged the activity of cell bodies (Methods). (F) Calcium response of an example SOM-neuron and a population of 178 SOM-neurons imaged in five mice. (G) The calcium response elicited by the background was stronger, on average, than that elicited by the figure (linear mixed-effects model, p<0.01). (H) Expression of GCaMP6f in V1 of an example PV-Cre mouse after AAV injection. We observed strong expression in the cell bodies. (I) Activity of an example PV-neuron and a population of 159 PV-neurons in four mice elicited by the figure-ground stimuli. (J) In PV-neurons, figures elicited stronger calcium responses than the background (linear mixed-effects model, p<0.001). (K) Significant FGM differences (difference in log ΔF/F between figure and ground) between neural classes (linear mixed-effects model: Bonferroni corrected post-hoc comparisons, ***, p<0.001). (L) To test a causal influence of VIP neurons in generating FGM, we optogenetically inhibited VIP neurons while recording V1 activity with electrophysiology. We injected AAV1-hSyn-SIO-stGtACR2-FusionRed in V1 of three VIP-Cre mice. (M) Average, normalized activity of a population of 168 recording sites (8 penetrations, 3 mice) elicited by an orientation-defined figure (dark purple) and background (light purple) without (left) and with (right) inhibition of VIP-neuron activity, starting 250ms after stimulus onset. (N) FGM, quantified using d-prime with (y-axis) and without (x-axis) optogenetic inhibition of VIP-neurons. Data of different penetrations are shown in distinct colors (linear mixed-effects model, p<0.05, time window 250-500ms).

To image the activity of VIP-neurons, we expressed GCaMP6f in VIP-Cre mice and observed that figures elicited stronger VIP-neuron activity than backgrounds (N=155 neurons in four mice; linear mixed-effects model; F_1,308_=9.1, p<0.01) (Figures 6B-D), consistent with the hypothesis that these neurons receive excitatory feedback from HVAs about salient visual stimuli (Zhang et al., 2014). To examine the activity of SOM-cells, we expressed GCaMP6f in SOM-Cre mice and found that activity elicited by figures was weaker than that elicited by the background (N=178 neurons in 5 mice, F_1,354_=7.6, p<0.01) (Figures 6E-G). This result supports the hypothesis that the release of SOM-inhibition contributes to the extra activity elicited by the figure (Karnani et al., 2016) (Figure 6A). We also examined the activity of parvalbumin positive (PV) cells, which have been implicated in feedforward inhibition and control of the gain of the cortical column (Harris and Shepherd, 2015; Ji et al., 2015). We expressed GCaMP6f in V1 of PV-Cre mice and observed that figures evoked stronger responses in PV-cells than backgrounds (N=159 neurons in 4 mice, F_1,316_=119, p<0.001) (Figures 6H-J). The level of FGM differed significantly between the different cell-types (linear mixed-effects model across cell-types, including the excitatory cells in Figure 4E; interaction between figure-ground condition and cell-type, F_1,3220_ = 32.5, p<0.001). FGM was strongest for PV cells, weaker and similar in magnitude in excitatory cells and VIP interneurons, and inverted in the SOM population (see Figure 6K for post-hoc comparisons of the different cell-classes). Visually evoked responses are modulated by locomotion (Dipoppa et al., 2018; Erisken et al., 2014; Niell and Stryker, 2010; Pakan et al., 2016). We observed that the activity of excitatory neurons, PV-, VIP- and SOM-positive interneurons were all enhanced by running (linear mixed-effects model; main effect of running, all ps<0.001), but there was no significant interaction between running and FGM in any of the cell types (p>0.05 for all cell-types).

### VIP-neurons contribute to FGM in V1

Do VIP-neurons indeed enhance the representation of figures in V1 as predicted by their proposed role in silencing SOM-neurons? To test the involvement of this disinhibitory circuit, we inhibited the activity of VIP-neurons and measured how it influences FGM in V1. We expressed an inhibitory opsin (stGtACR2) in VIP-neurons in V1 and suppressed their activity, while electrophysiologically recording V1 activity evoked by orientation-defined figures and the background (Figures 6L, S7E). We started the inhibition of VIP-neurons 250ms after stimulus onset, to selectively influence the late response phase during which HVAs feed back to V1 and to prevent interference with the early phase during which V1 neurons propagate activity to HVAs. We found that inhibiting VIP-activity reduced FGM in V1, decreasing the d-prime by 47% (Figures 6M,N linear mixed-effects model, p<0.05, 3 mice, 8 penetrations, 168 recording sites, time-window 250-500ms). This finding supports the role of VIP-neurons in generating FGM in V1.

## Discussion

Our results provide new insights into how the interactions between V1 and HVAs shape perception. Upon appearance of a new image, the activity of V1 neurons exhibits a number of phases (Lamme and Roelfsema, 2000). During an early feedforward phase, activity is propagated from the retina via the thalamus to V1 and is transmitted onwards to HVAs. V1 activity during this phase suffices for ceiling performance in the relatively simple contrast detection task, as if V1 acts as relay station that only needs to transmit information about the stimulus to HVAs (Thorpe et al., 1996). Inhibition of the entire visually driven V1 response reduced the accuracy of the mice. In accordance with a previous study (Prusky and Douglas, 2004), most animals were still able to perform the task above chance level, suggesting that information about the visual stimulus still reached the brain regions that are required for an appropriate licking response via alternative routes. Although we do not know what the mice perceived during V1 inhibition, comparable effects have been reported in humans with lesions in V1, who are able to correctly guess the presence, location and shape of simple visual stimuli in the affected area of the visual field, despite denying any visual awareness of the stimuli (Pöppel et al., 1973; Sanders et al., 1974; Weiskrantz et al., 1974). This phenomenon is known as blindsight and has been replicated in monkeys with a V1 lesion (Cowey and Stoerig, 1995; Cowey and Weiskrantz, 1963), where information from the LGN is directly propagated to HVAs, bypassing V1 (Schmid et al., 2010).

If a visual stimulus requires figure-ground segregation, however, the peak response is followed by a phase in which the V1 neurons exhibit FGM; the representation of figural image elements is enhanced. In previous work, it remained unclear whether this late V1 activity phase is useful for perception, but our results demonstrate that optogenetic V1 silencing during this phase blocks figure-ground perception. This finding is in line with previous TMS studies in humans, although TMS produces much weaker perceptual effects (Wokke et al., 2012). Among the studied figure-ground stimuli, we observed differences in the latency and strength of FGM, measured electrophysiologically (Figure 2B), and in the V1 processing time required for figure-ground perception. FGM of orientation-defined figures occurred earlier and was stronger than FGM of phase-defined figures and the mRT was also shorter for orientation-defined stimuli. Accordingly, V1 inhibition interfered with figure-ground perception at a later point in time for phase-defined than for orientation-defined figures.

The present results seem to differ from previous studies that examined the early and late phases of neuronal responses in the barrel cortex of mice engaged in tactile detection tasks (Manita et al., 2015; Sachidhanandam et al., 2013). Suppression of the late activity in the barrel cortex interfered with the detection of a tactile stimulus whereas in the present study late V1 activity was not required for the contrast detection task. We speculate that this difference between results may be related to the use of relatively weak tactile stimuli in these previous studies, which may have required amplification by recurrent interactions between lower and higher areas of the somatosensory cortex.

We identified a source for V1-FGM in HVAs that also exhibit FGM and project back to V1. A number of HVAs exhibited FGM and it was strong in areas lateral from V1 (LM and AL) during passive viewing and weaker in anteromedial areas (RL, AM and PM), although the wide-field experiments demonstrated that FGM increases in these areas when mice use the figure-ground stimuli to perform a task. Understanding the functional organization of the mouse visual cortex is an active field of exploration (Juavinett and Callaway, 2015; Marshel et al., 2011; Smith et al., 2017; Wang et al., 2011, 2012), and the present results suggest that figure-ground segregation can help to further dissect the roles of higher cortical areas.

Optogenetic silencing of the HVAs strongly reduced FGM in V1, without a consistent effect on the early, visually driven response. This finding supports theories suggesting that FGM in V1 requires feedback from HVAs (Bhatt et al., 2007; Roelfsema et al., 2002). The effect of silencing was pronounced for the phase- and texture-defined figure-ground stimuli, and somewhat weaker if the figure was a grating with a different orientation. The weaker suppression is compatible the view that horizontal interactions within V1 may also contribute to the perception of orientation-defined figures. Horizontal interactions within V1 are thought to be orientation selective so that suppression is stronger within image regions with a homogenous orientation, and weaker if a figure with one orientation is superimposed on a background with another orientation (Li, 1999). If the figure is defined by a phase offset, the orientation selective suppression signal is absent, which may explain the larger dependence on cortico-cortical feedback. We note, however, that the viral construct was not taken up by all neurons in HVAs so that inhibition was presumably incomplete. Hence, we cannot exclude that the remaining FGM for orientation-defined figures reflects a feedback influence from the non-silenced neurons in HVAs.

The main effect of silencing activity in HVAs was a decrease in activity during the late phase of the V1-response, which was more pronounced for figures than the background, thereby decreasing FGM (Figure S6). Importantly, these excitatory feedback effects imply that FGM is not caused by surround suppression and are consistent with predominantly excitatory (or disinhibitory) feedback effects on V1 activity (Huh et al., 2018; Hupé et al., 1998; Keller et al., 2020). However, a study in monkeys (Nassi et al., 2013) demonstrated that the cooling of HVAs increased V1 activity evoked by large stimuli, which suggested a suppressive influence of feedback connections. We do not know if this discrepancy reflects a difference between species or between methods used for silencing.

A previous study (Zhang et al., 2014) suggested that feedback connections enhance V1 activity at relevant locations by activating VIP-interneurons, which in turn inhibit inhibitory SOM-cells, thereby disinhibiting the cortical column. In accordance with this hypothesis, we observed that figures enhanced the activity of VIP-neurons and suppressed the activity of SOM-cells (Figure 6A), which supports the view that SOM-interneurons suppress the V1 activity elicited by homogeneous image regions (Adesnik et al., 2012). Inactivation of VIP-neurons decreased V1 activity and diminished FGM, demonstrating that VIP-mediated disinhibition contributes to the enhanced activity elicited by figures.

PV-neurons were more active for figures than for the background, and this FGM-signal was even stronger than that of pyramidal cells. Although the PV-activity profile resembles that of VIP-neurons, PV-cells do not preferentially contact other inhibitory cell types to cause disinhibition. Instead, they suppress pyramidal neurons (Atallah et al., 2012; Pfeffer et al., 2013; Williams and Holtmaat, 2019), making it unlikely that they are part of a disinhibitory circuit motif. PV-cell activity was weak for the background, which, unlike for SOM-cells, also rules out a specific role for PV-neurons in suppressing the pyramidal cell activity elicited by the background. Instead, our results support the hypothesis that PV-neurons integrate the activity of nearby pyramidal neurons to control the gain of the cortical column. If pyramidal neurons become more active, the activity of nearby PV-cells also increases and their input suppresses pyramidal neurons, giving rise to a negative feedback loop (Figure 6A). The high PV-cell activity level elicited by figures is also in accordance with a study in the visual cortex in monkeys (Mitchell et al., 2007), which demonstrated that fast-spiking interneurons strongly increase their activity for behaviorally relevant stimuli.

The present results provide insight into how interactions between lower and higher areas of the visual cortex enable the co-selection of image elements that belong to a single figure and their segregation from the background. These interactions are in part mediated by direct corticocortical connections, but there are also routes through subcortical structures (e.g. LP of the thalamus (Sherman, 2016)) and future studies could compare the relative contributions of these different routes. As a result of these interactions, figural image elements become labeled with enhanced neuronal activity across multiple areas of the visual cortex, in accordance with the view that such a labeling process determines the perceptual grouping of features of the same object (Duncan et al., 1997; O’Craven et al., 1999; Roelfsema, 2006; Roelfsema and Houtkamp, 2011). It seems likely that the findings generalize to more complex forms of perceptual organization. Imagine grasping an object that is surrounded by a number of other objects. The visual system guides our fingers to touch and grasp edges of the same object, a selection process (Baldauf and Deubel, 2010) that has to rely on recurrent interactions between higher areas coding for the relevant object’s shape and lower areas coding for its individual edges. Previous studies in monkeys demonstrated that the representation of a selected object’s shape is enhanced in higher cortical areas (Chelazzi et al., 1993) and that this also holds true for relevant edges in V1 (van Kerkoerle et al., 2017; Roelfsema and Houtkamp, 2011). The present study illustrates how such selection process is coordinated across cortical areas. Future studies can now start to investigate the interactions between brain regions that are required for complex forms of perceptual organization and how they guide behavior (Roelfsema, 2006).

## Acknowledgments

We thank the animal and mechatronics department at the Netherlands Institute for Neuroscience and Isis Alonso, Konstantinos Koukoutselos, Lola van Linge, Tobias van der Bijl, Veerle Bos and Anthony Poerwoatmodjo for their assistance in mouse training and electrophysiological recordings. We thank Emma Ruimschotel for support with breeding and Leonie Cazemier for helpful comments. The work was supported by NWO (ALW grant 823-02-010) and the European Union’s Horizon 2020 and FP7 Research and Innovation Program (grant agreements 720270 and 785907 ‘‘Human Brain Project SGA1 and SGA2’’, ERC grant agreement 339490 ‘‘Cortic_al_gorithms’’, FLAG-ERA JTC grant ChampMouse and the Erasmus Mundus “NeuroTime” program) and the Friends Foundation of the Netherlands Institute for Neuroscience.

## Author contributions

UHS, EvB and JAML developed the task. LK and MWS recorded and analyzed the electrophysiological data in which mice were passively viewing the stimuli. LK performed the V1 optogenetic silencing experiments during task performance. EvB and AB performed the wide-field experiments and EvB and MWS analyzed the results. SM carried out the two-photon experiments and MWS, SM, UHS and CvdT analyzed the two-photon data. LK and MWS recorded and analyzed the electrophysiological data in which we inhibited VIP-neurons and the recordings in which we silenced higher visual areas. UHS performed and analyzed the electrophysiological experiment during task performance. CL helped with the design of experiments involving transgenic mouse lines. PRR and MWS conceived of the experiments based on advice from JAH and supervised the project. PRR, MWS and LK wrote the paper.

## Data and materials availability

All data and the computer code used to analyze the data will be made available for download and curated at the Human Brain Project Joint Platform. Correspondence and requests for materials can be sent to PRR.

## STAR Methods

### Contact for Reagent and Resource Sharing

Correspondence and requests for materials can be sent to PRR.

### Experimental Model and Subject Details

35 male and 14 female mice of 2-14 months age were used in this study. All experimental procedures complied with the National Institutes of Health Guide for Care and Use of Laboratory Animals and the protocol was approved by the ethical committee of the Royal Netherlands Academy of Arts and Sciences and the CCD. The experiments were performed in accordance with the relevant guidelines and regulations.

### Method Details & Quantification and Statistical Analysis

#### Visual Stimuli

We created the visual stimuli with the Cogent toolbox (developed by John Romaya at the LON at the Wellcome Department of Imaging Neuroscience) and linearized the luminance profile of the monitor/projector. Visual stimuli during passive electrophysiological experiments were projected onto a back-projection screen placed 15cm from the mouse with a PLUS U2-X1130 DLP projector (mean luminance = 40.6cd.m^-2^). The size of the projection was 76×56cm, the field-of-view 136° x 101.6°, the resolution 1024×768 pixels and the refresh rate 60Hz. For the behavioral electrophysiological experiments, the stimuli were presented on a 21-inch LCD screen with a resolution of 1280×720 pixels (Dell 059DJP) driven by a Windows computer at 60Hz at a distance of 15cm in front of the mouse. During the two-photon experiments, we presented visual stimuli to the left eye of the mice, using a 24-inch LCD monitor (Dell U2414H) with a resolution of 1920×1080 pixels and a refresh rate of 60Hz, placed at an angle of 30° relative to the nose and a distance of 12cm from the eye. For the optogenetics experiments, we used a 24-inch LCD monitor (1920 x 1200 pixels, Dell U2412M), placed 11cm in front of the eyes and for wide-field experiments an LCD monitor (122 x 68cm, Iiyama LE5564S-B1), at a distance of 14cm. We applied a previously described (Marshel et al., 2011) correction for the larger distance between the screen and the mouse at higher eccentricities. This method defines stimuli on a sphere and calculates the projection onto a flat surface. The orientation- and phase-defined figure-ground stimuli were composed of 100% contrast sinusoidal gratings with a spatial frequency of 0.075 cycles/deg. and a mean luminance of 20cd/m^2^. The diameter of the figure was either 35° (optogenetic- and wide-field imaging experiments), 40° (electrophysiology during passive viewing), 50° (two-photon experiments). For the orientation-defined figures (Figures 1A,B), the grating orientation in the background was either horizontal or vertical and the orientation of the figure was orthogonal. For the phase defined figures (Figure 1A), the phase of the figure grating was shifted by 180° relative to that of the background. For the contrast-defined stimuli, we presented the figure gratings on a grey background (20cd/m^2^). To test the generality of figure-ground perception, we also presented random textures in some of the experiments, again ensuring that the figure and background stimulus in the RF of the neurons was identical. The texture was made by filtering Gaussian distributed random noise patterns through an oriented filter. Four new random noise patterns were generated for each experimental session and these were filtered with 0° and 90° filters, yielding eight oriented textures. The oriented filter (F) was made through summation of individual Gabor filters (G) as follows:

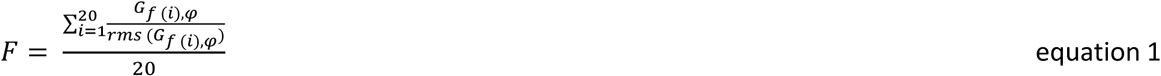

Where G is Gabor filter of spatial frequency *f (i)*, where *f* was linearly spaced from 0.05 to 0.5 cycles/deg, orientation φ (0° or 90°) and standard deviation *1/f, rms* indicates a root-mean square operation. After filtering, the texture was matched to the root-mean square contrast of the sine-wave grating used in the other experiments using an iterative hard-clipping procedure with 10 iterations.

#### Anesthesia during surgeries

The mice were handled five to ten minutes per day starting one week before surgery. Anesthesia was induced using 3-5% isoflurane in an induction box and it was maintained using 1.2-2.5% isoflurane in an oxygen-enriched air (50% air, 50% O_2_) mixture and we subcutaneously injected 5 mg/kg meloxicam (0.5 mg/ml) as general analgesic. The mice were positioned in a stereotactic frame and we monitored the depth of anesthesia by frequently checking paw reflexes and breathing rate. The temperature of the animal was monitored and kept between 36.5° and 37.5° with a heating pad coupled to a rectal thermometer. We covered the eyes with ointment to prevent dehydration. The area of incision was shaved, cleaned with betadine and lidocaine spray was applied to the skin as a local analgesic.

#### Behavioral Task

The mice were held on a reverse day-night cycle and a fluid restriction protocol with a minimal intake of 0.025ml/g, while their health was carefully monitored. The animals were trained to indicate the side on which a figure appeared by licking the corresponding side of a custom-made double lick spout (Figure 1B). We registered licks by measuring a change in capacitance with an Arduino and custom-written software. A trial started when the stimulus with a figure on the left or right appeared on the screen. The stimulus was displayed for 1.5 seconds. Because mice made early random licks, we disregarded licks from 0-200ms (grace period). We prolonged the grace period to 500ms for one mouse, as it helped correct a bias for preferentially licking one side (mouse M2 in Table S1). The exact figure location varied slightly depending on the experiment, but the figure center was generally close to an azimuth of ±30° (left or right of the mouse) and an elevation of 15°.

Stimulus presentation was followed by a variable inter-trial interval (ITI) of 6-8s. Correct responses were rewarded with a drop of water or milk. If the animal made an error, a 5s timeout was added to the ITI. We presented a background texture during the ITI and did not give reward if the mice licked so that they learned to ignore it. In some sessions, we included correction trials, which were repeats of the same trial after an error. We only included the non-correction trials to compute accuracy, defined as hits/ (hits+errors). During task performance in the wide-field imaging experiments, we used a motor which moved the lick spout close to the mouth of the mouse, 500ms after the presentation of the stimulus, thereby enforcing a minimum viewing time before the mouse could respond. For the accuracies of individual mice see Table S1.

To train mice on the figure-ground task, we first trained them to detect grating figures on a grey background. The background was then replaced by a grating with a gradually increasing contrast across several weeks of training.

#### Electrophysiology

The laminar electrophysiological recordings during passive viewing (Figures 2A,B) were carried out in six Tg (Thy1-GCaMP6f)GP5.17Dkim mice (5 male, 1 female) aged between two and six months. After induction of general anesthesia (as described above), the skin on the skull was opened, tissue was removed from the skull and we briefly applied H_2_O_2_. We then applied a dental primer (Keerhawe Optibond) to improve the bonding of cement to the skull. We applied a thin layer of dental cement (Heraeus Charisma A1) across an area of the skull anterior to bregma and fixed a head bar with additional dental cement (Vivadent Tetric Evoflow). We implanted one or two small screws over the cerebellum, which served as reference and ground wires. We used the dental cement to fix them in place and build a well around the skull covering the posterior part of the left hemisphere, to prevent growth of the skin over the area of interest. After three weeks, we made a craniotomy centered on the area of V1 with a population RF at 30° azimuth and 20° elevation. The mice were head-fixed and placed on a treadmill so that they could run or sit according to their preference. We tracked the treadmill movements using an Arduino and monitored pupil movements and size under infrared light with a zoom lens (M118-FM50, Tamron, Cologne, Germany) coupled to a camera (DALSA GENIE-HM640, Stemmer Imaging) and custom written software. We inserted a linear-array recording electrode (A1×32-5mm-25-177, NeuroNexus, 32 channel probe, 25 micron spacing) in V1 and lowered it to around 1mm below the brain surface and adjusted the depth of the electrode with reference to the current source density profile as reported previously (Self et al., 2014) to ensure coverage of all layers. We amplified the electrical signal from the electrodes and sampled it at 24.4kHz using a Tucker-Davis-Technologies recording system. We removed muscle artifacts by re-referencing each channel to the average of all other channels before filtering the signal between 500 and 5000Hz. We detected spikes by thresholding (positive and negative threshold) the band-passed signal at 4 times an estimate of the median absolute deviation and convolved the detected spikes with a Gaussian with a standard deviation of 1.3ms (and an integral of 1) to derive an estimate of multi-unit spike-rate. First, we measured the RF of the units recorded at each electrode with a sparse noise stimulus consisting of 4 white checks (8 by 8°, 40cd/m^2^) on a black background presented for 250ms with a 250ms inter-trial interval. The checks (>30 presentations per check) were positioned on a grid ranging from -64 to 16° horizontally and -22 to 66° vertically relative to the mouse’s nose with negative values indicating the right hemifield. We corrected for flat screen distortion as was described above. We averaged the MUA response evoked by each check in a time window from 50-400ms after check onset to obtain a map of visual responsiveness and fit a 2D-Gaussian to estimate the width and center of the RF (Figure 2A). The quality of the fit was assessed using r^2^ and a bootstrapped variability index (BVI), which estimated the reliability of the RF center estimate. We resampled an equal number of trials as in the original dataset (with replacement) and regenerated the Gaussian fit. The BVI is defined as the ratio of the standard deviation of the RF center position and the standard deviation of the fitted Gaussian. To measure FGM, we centered a 40° diameter figure on the RFs of layer 4 units. To create the background condition, we shifted the figure by 50-60° so that the RF fell on the background. As the RF position varies slightly across layers we quantified for each recording site the percentage of the RF area that fell within the figure boundary. We only included recording sites if (i) the overlap between the RF and the figure was greater than 70% to exclude boundary driven responses, (ii) the RF was reliable (r^2^ of the Gaussian fit > 0.33, BVI < 1.5), (iii) if the signal-to-noise ratio (SNR) of the visual response was greater than 1 (ratio of the activity between 0-100ms after stimulus onset to the standard deviation of baseline activity [-200-0ms] across trials) and (iv) if the maximum response of the site was greater than 2Hz. These criteria led to the inclusion of 198 recording sites for the electrophysiological data (Figure 2B). The orientation-, phase- or texture-defined figure-ground stimuli were presented in blocks of 32 stimuli. The position of the figure and the orientation of the underlying pattern/texture was pseudorandomized within the block so that the RF-stimulus was identical for the figure and ground conditions. To generate population responses, we subtracted the pre-stimulus activity (time-window -200-0ms relative to stimulus onset) and normalized the activity at each recording site to the peak of the average, smoothed (lowess method, 39ms window size) response across the figure and ground conditions (time-window, 0-100ms).

FGM was quantified with the d-prime, which is a measure for the reliability of the signal on individual trials (time window 100-500ms):

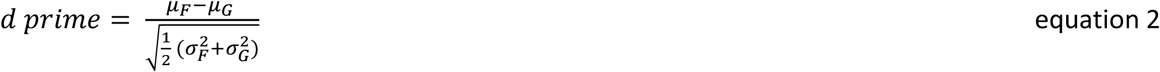

Where *µ*_*F*_ and *µ*_*G*_ are the means and 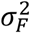 and 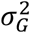 are the variances of the figure and ground response across trials, respectively. To estimate whether the level of FGM, as measured with d-prime, differed between different cue-types (i.e. orientation/phase/texture) we modelled the hierarchical correlation structure in the data with linear mixed-effects models to assess significance (Scheffe, 1956) using the fitlme.m function in Matlab, because every electrode contained multiple contacts and all contacts were tested for each of the cue types. Hence, there were two random effects: the recording contacts and the penetrations, which were included as random intercepts in the model. The two fixed effects were cue-type (orientation-defined, phase-defined and texture defined figure) and the overall intercept (i.e. mean d-prime for the reference condition, which was one of the cue-types). We obtained best model fits (as judged by a lower Akaike’s Information Criteria (AIC)) if we included a random slope term for cue-type grouped by penetration. Differences between cue-types were assessed by Bonferroni corrected post-hoc contrasts. To assess whether FGM was significant for each cue, we fit three separate linear mixed-effects models with a fixed intercept term and a random effect term for the penetration. As in previous studies, we estimated the latency of FGM by fitting a function (Poort et al., 2012) to the figure minus background response in a time-window from 0 to 300ms after stimulus onset. In brief, the function is the sum of an exponentially modulated Gaussian and a cumulative Gaussian, capturing the Gaussian onset of neural modulation across trials/neurons and the dissipation of modulation over time. The latency (small arrows in Figure 2B) was defined as the (arbitrary) point in time at which the fitted function reached 33% of its maximum value. Statistical comparisons of latency estimates were performed by bootstrapping. We selected a random set of recording sites (with replacement) 1000 times and recalculated the latency estimate for each resampled population.

#### Optogenetic silencing of V1 during contrast and figure-ground perception

For optogenetic silencing of V1 during behavior (Figures 2C-E) we injected five C57BL/6 mice (4 male, 1 female) aged between 2 and 14 months with a cell-specific viral vector (AAV5-CaMKII-hGtACR2-eYFP) coding for the inhibitory GtACR2 opsin (Govorunova et al., 2015). The virus was cloned and produced in a custom preparation by Virovek, Inc., Hayward, CA, USA based on the plasmid pFUGW-hGtACR2-EYFP, which was a gift from John Spudich (Addgene plasmid #67877; http://n2t.net/addgene:67877; RRID:Addgene_67877). The general surgical procedures were described above. For the viral injections, an incision in the skin was made and the skin was gently pulled aside, exposing the area of the skull above the cortex. We used a dental drill to make small craniotomies above V1 of both hemispheres 2.7mm lateral from the midline, 0.5mm anterior of lambda. We placed a pulled borosilicate capillary containing the virus vertically above the craniotomy touching the dura. Slowly the pipette was lowered to a depth of 600µm from the brain surface and we slowly injected a total of 150nL per hemisphere with a concentration of 3*10^11^ GC/mL, at different depths using Nanoject III programmable injector (Drummond Scientific). After the injection, we left the pipette in place for at least 8 minutes before slowly retracting it to avoid efflux of virus. After two weeks, we attached a head-fixation bar to the skull (see electrophysiological procedures). We applied a thin layer of adhesive to the bone which clarifies the skull, a method referred to as the ‘clear skull cap’ technique (Guo et al., 2014). After the adhesive dried, we applied a thin layer of transparent dental cement (C&B Super-Bond) on top. We added a small rim of dental cement to the outer edges of the skull cap to prevent growth of the skin over the area of interest. Before we started the optogenetic silencing experiments, we shortly anesthetized the mice with isoflurane (see above) to apply a thin layer of transparent nail polish (Electron Microscopy Sciences) to the cap to reduce light glare.

Once the animals performed the task consistently with an accuracy larger than 65%, we introduced trials in which neural activity in the visual cortex was inhibited by activating the opsin. We presented figure-ground stimuli (defined by a difference in orientation or phase) on 75% of all trials and applied optogenetic silencing in a random 25% of those trials. We presented contrast-defined stimuli in 25% of all trials and because the accuracy was high and stable, we increased the fraction of trials with optogenetic intervention up to 50% (we used the same trial ratios in control experiments in which the laser was not directed at V1; Figures S3D,E). We used a DPSS Laser (BL473T3-100FC, Shanghai Lasers & Optics Century Co.) emitting blue light (wavelength 473nm) as a light source and directed the light to the cortex with a split optical fiber (2×2 Coupler, 50:50, 200µm, FC/PC to 1.25mm Ceramic Ferrule, Thorlabs, Newton). The two fiber ends were directed at the centers of area V1 in the two hemispheres. Optogenetic stimulation lasted for 2s with a constant power of 10mW at each fiber end. The laser power at the skull cap was 5.6mW/mm^2^. The clear skull cap absorbs around 50% of the light (Guo et al., 2014) so that the effective laser power at the cortical surface was ∼2.8mW/mm^2^, a light level that does not cause measurable heating (Owen et al., 2019). The onset of stimulation was shifted relative to the onset of the visual stimulus in steps of 16.7ms according to the frame rate of the screen. The time between stimulus onset and laser onset was 17, 33, 50, 67, 83, 100, 150 or 200ms. To remove spurious cues that might be caused by optogenetic stimulation, we placed a light shield around the head of the mouse to prevent light from the optogenetic stimulation reaching the eye and presented blue light flashes at random times by driving a blue LED placed below the mouse’s head to flash every 0.5-1s with an Arduino in all optogenetic experiments. We only included sessions in which the overall accuracy of the mice on laser off trials was above 70%. We used correction trials, which were repeats of the same condition after an error, to decrease a response bias and excluded these trials when computing accuracy (correction trials were not used for optogenetic silencing trials). We used binomial tests (with Bonferroni-correction) to examine whether performance was significantly above chance level during trials with optogenetic manipulation and to test whether optogenetic inhibition impairs performance. To estimate the time at which the accuracy reaches its half maximum, we fit a logistic function to the accuracy as a function of the laser onset latency using the Palamedes toolbox in Matlab. We used bootstrapping (1000 times) by sampling trials with replacement to determine the 95%-confidence interval of the latency, defined as the time when accuracy was halfway between the earliest and latest V1 silencing time point (i.e. the inflection point of the fitted function).

#### Wide-field imaging

In the wide-field imaging experiments (Figure 3) we included 8 male and 1 female Thy1-5.17 GCaMP6f mice (Dana et al., 2014) aged between 2 and 14 months. The skull of the animals was prepared with the clear-skull cap technique (described above). We placed the mice under a wide-field fluorescence microscope (Axio Zoom.V16 Zeiss/Caenotec) to image a large part of the cortical surface. Images were captured at 20Hz by a high-speed sCMOS camera (pco.edge 5.5) and recorded using the Encephalos software package (Caenotec). We monitored the size and position of the right pupil (100Hz sampling rate) and movements of the mouse with a piezo plate under the front paws of the mouse (100Hz sampling rate) and removed trials with large body movements in a time-window from the start of a trial until 350ms after stimulus onset. We captured images of 1600×1600 pixels (∼15µm per pixel), down-sampled them to 400×400 pixels and applied a Gaussian filter smoothing kernel of 5×5 pixels. We carried out a population RF mapping session (see below) to determine the borders of V1 and the HVAs and we matched these areas to the Allen Brain common coordinate framework (Figure S5D). We computed the average ΔF/F (relative to the baseline fluorescence in a 300ms window before stimulus onset) of all pixels within an area. In the active task, trials were removed if (i) the mouse had a performance below 65% in a window of fifteen trials, or (ii) the absolute z-score of ΔF/F was larger than 3.5 (removal of trials with artifacts). We measured FGM with the d-prime in a time window of 150-300ms after stimulus onset, averaged across both hemispheres. We assessed significance with repeated measures ANOVAs, testing whether the d-prime was higher than 0. In addition, we used a repeated-measures ANOVA to test the effect of area (i.e. V1, LM, AL, RL, A, AM, PM, RSP, M1 and M2) and response type (hit, error or passive viewing). We carried out the passive viewing experiments before the start of training in the figure-ground task (N=5). See Tables S1, S2 for information about the mice in this experiment.

#### Two-photon imaging of excitatory neurons

We carried out experiments with five Thy1 mice to image the activity of excitatory neurons (Figure 4). All mice were between 2-6 months old and included both sexes (Table S2). The animals were anesthetized as described above. Mice were first implanted with a head-ring for head-fixation and after two weeks of recovery, we mapped the retinotopy (see below for the pRF mapping method) to determine the boundaries of visual areas. These mice were additionally injected with AAV1-CaMKII-GCaMP6f-WPRE-SV40 (Penn Vector Core, University of Pennsylvania, USA) in V1 (100nl) and LM, AL, RL, AM, PM (50nl each) at an injection speed of 20nl/min distributed across two depths (400µm, 200µm below the pial surface) to enhance the GCaMP signal. All viral titers were adjusted to 10^12^GC/ml before injection. We made a circular craniotomy with a diameter of 4-5mm centered at 0.5mm anterior to lambda and 2.5 lateral from the midline. We carefully thinned the bone along the outer diameter of the craniotomy and slowly lifted the bone flap without damaging the dura, which was kept moist with warm ACSF or saline. The craniotomy was closed with a double-layered glass coverslip, with the outer glass resting on the skull, which was fixed with dental cement (Vivadent Tetric Evoflow). After two weeks, we habituated the mice to head immobilization while they could run on a running belt under the two-photon microscope (Neurolabware). We imaged through a 16x water immersion objective (Nikon, NA 0.80) at 1.7x zoom at a depth of 120-300µm with a 15.7Hz frame rate and a resolution of 512 x 764 pixels. We targeted V1 and other HVAs based on the retinotopic maps. A Ti-Sapphire femtosecond pulsed laser (MaiTai, Spectra Physics) was tuned to 920nm for delivering excitation light. The power used varied between 20 and 50mW depending upon the depth of the imaging plane. First, we mapped the RF locations of the neurons within the field of view. We presented 12 x 12° white (38cd/m^2^) squares on a black background (0.05cd/m^2^) in an area ranging from -18 to 78° horizontally and -21 to 51° vertically relative to the mouse’s nose. Each square was flashed twice for 166ms with a blank interval of 166ms, followed by delay of 500ms. The square at each location was presented 20 times. We calculated RFs based on the response in a 500ms window after stimulus onset. We fit a linear regression model to estimate the responses to the squares of the grid, regressing out the influence of running and the interaction between the visual stimulus and running. We fit a circular 2D-Gaussian to the beta values for every location to estimate the RF center and its full width at half-maximum response strength. We evaluated the quality of the fit using the r^2^ value and the BVI (see above; r^2^ of the Gaussian fit > 0.33, BVI < 1.5). The figure-ground stimuli contained a 50° figure with one of 6 different orientations (maximum luminance of 38cd/m^2^ and minimum luminance of 0.05cd/m^2^). The figure was presented 120 times at two positions on the screen (20 repetitions per orientation), one centered on the RF location of a cluster of imaged cells, and the other at a distance of 55° from the RF center. The stimuli were presented in randomized order in blocks of 48 trials for 0.5s with an ITI of 2.5s to allow enough time for decay of the calcium signal of the previous trial. We ensured that the stimulus in the RF was identical, on average, in the figure and background conditions, as described in the main text. We used CAIMAN (Pnevmatikakis and Giovannucci, 2017; Pnevmatikakis et al., 2016) for pre-processing. We performed rigid motion correction for small shifts in the data due to motion of the animal, followed by the extraction of ROIs and the ΔF/F. ROI components found by the CNMF algorithm (Pnevmatikakis et al., 2016) were classified into neuronal compartments and noise using a Keras pre-trained convolutional neural network (CNN) classifier (CAIMAN matlab github library; https://github.com/flatironinstitute/ CaImAn-MATLAB). We only included ROIs that belonged to cell bodies in the analysis. To generate population responses, we normalized the activity of each cell by subtracting base-line activity (time-window -325-0ms [5 frames] before stimulus onset) and dividing by the maximum of the average of the figure and ground conditions (time-window 0-2s). For statistical analysis, we took the average baseline corrected ΔF/F values in a window from 0.3-1.5s after stimulus onset for each cell in the figure and ground conditions. The distribution of these values was positively skewed and we therefore took the log transform of the average ΔF/F values. Because the logarithm of negative values is undefined we first removed cells with negative responses in either the figure or ground conditions. We removed outliers using an iterative multi-variate outlier removal process. The Mahalanobis distance of each cell to the mean of the full distribution was calculated and z-scored. We removed cells for which the absolute z-score was greater than 3. If the maximum value of the pre-removal z-score was greater than 6 (indicating the presence of an extreme outlier which may distort the calculation of the z-score) the process was repeated. This procedure removed less than 3% of cells. This outlier removal method was performed once for all the excitatory cell data from the different visual areas, after concatenating the data from each area. Similarly, the data from the different interneuron sub-classes (described below) were concatenated together with the data from the excitatory cells in V1 before applying the outlier removal algorithm. The significance of the differences between figure and ground were first assessed using an omnibus linear mixed-effects model. Two models were analyzed, one for the data from the excitatory cells in different visual areas (i.e. the data shown in Figure 4) and one for the data from different interneuron sub-classes (including the excitatory cells in V1, i.e. the data in Figure 6). The models contained two within-cell fixed effects terms (intercept [the mean log (ΔF/F) in the background] and figure-ground condition) and one between-cell factor (either ‘Visual-area’ or ‘Cell-class’ depending on the model). The models also contained two random intercept terms for the cell identity and the imaging session in which the cell was recorded to account for any increased co-variance between cells from the same imaging session. We also ran post-hoc models for each visual area and cell-class separately. These models contained two fixed effects (intercept and figure-ground condition) and two random intercept terms (cell identity and imaging session).

#### Electrophysiology with optogenetic inhibition of higher visual areas

The laminar electrophysiological recordings with optogenetic inhibition of HVAs (Figure 5) were carried out in five male Tg (Thy1-GCaMP6f)GP5.17Dkim mice aged between two and six months. Surgical procedures were identical to those described above (under electrophysiology). We additionally applied a thin layer of clarifying adhesive to the skull of the left hemisphere (clear skull cap (Guo et al., 2014)). After one week of recovery, we used population RF mapping based on the GCaMPf expression (see pRF mapping below) to determine the borders between the visual areas. We targeted virus injections to HVAs based on these borders. We anesthetized the animals as described above and made four small craniotomies, each at a minimum distance of 500 µm from the V1 border. We slowly lowered a pulled borosilicate capillary containing the virus (AAV1-CaMKII-stGtACR2-FusionRed, titer 1.5×10^13^) to a depth of 600µm from the brain surface and slowly injected a total of 40nL per injection site (15nL at 600µm, 10nL at 400µm, 15nL at 200µm) using a Nanoject III programmable injector (Drummond Scientific). The construct encoding soma-targeted GtACR2 (Mahn et al., 2018) (pAAV-CKIIa-stGtACR2-FusionRed) was a gift from Ofer Yizhar (Addgene viral prep #105669-AAV1; http://n2t.net/addgene:105669; RRID:Addgene_105669). After the injection, we left the pipette in place for five minutes before slowly retracting it to avoid efflux of virus. We sealed the chamber with the biocompatible adhesive Kwik-Cast (WPI). After three weeks, we made a craniotomy centered on the area of V1 with a population RF at 30° azimuth and 20° elevation. The mice were head-fixed on a treadmill and we performed laminar recordings, RF mappings and placed figures as described above (under electrophysiology). We activated the laser on 50% of the trials, randomly interleaved with the trials without optogenetic inactivation. We used a DPSS Laser (BL473T3-100FC, Shanghai Lasers & Optics Century Co.) emitting blue light (wavelength 473nm) with a constant power of 5mW as light source and directed the light to lateral and medial HVAs through a split optical fiber (2×2 Coupler, 50:50, 200µm thickness, FC/PC to 1.25mm Ceramic Ferrule, Thorlabs, Newton). The laser was turned on 200ms before onset of the visual stimulus and remained on for the entire duration of visual stimulation (500ms) and was turned off 100ms after stimulus offset. We normalized V1 activity to that in the figure and ground conditions without optogenetic intervention. We quantified FGM using d-prime as described above and removed extreme multi-variate outliers (likely artifacts) by calculating the z-scored Mahalanobis distance of each recording site’s d-prime_laser off_ and d-prime_laser on_ values from the full distribution. Recording sites with values of greater than 2.58 were removed (approximately 1% of recording sites). To estimate the significance of the laser-induced change in d-prime we fit the data with a linear mixed-effects model containing two fixed effects (laser on/off, and intercept, i.e. d-prime when the laser was off), two random intercept terms to account for co-variability of data obtained from the same electrode contact site and penetration and one random-slope term to account for the fact that the effect of the laser varied across penetrations (Figure S6D). The same model was used to assess the laser-induced change in peak response (mean normalized activity in a window from 0-100ms after stimulus onset). Laser-induced changes in figure and ground activity were assessed with a post-hoc test including a random intercept term for electrode penetration.

#### Two-photon imaging of inhibitory neurons

We carried out experiments with four VIP-Cre, five SOM-Cre and four PV-Cre animals between 2-6 months old including both sexes (Figures 6A-K, Table S2). The animals were anesthetized as described above. An incision in the skin was made along the anteroposterior axis, and the skin was gently pulled to the side, exposing the area of the skull above the cortex and the area posterior to lambda. We drilled a small craniotomy over the center of right V1 (AP 0.5mm anterior to lambda and 2.5mm lateral from the midline) and slowly injected 200-300nL of the virus (AAV9-CAG-flex-GCaMP6f-WPRE-SV40, Penn Vector Core, University of Pennsylvania, USA) at 20nl/min distributed across three depths (500µm, 300µm and 150µm from the pial surface). The craniotomy was sealed and the skin was sutured. Two weeks later, the animals underwent a second surgery to implant a head-ring for immobilization and a cranial window to allow imaging of neuronal activity (see description of two-photon imaging method above). After two weeks, we habituated the mice to head immobilization while they could run on a running belt under the two-photon microscope. We mapped the RF of the neurons, presented figure-ground displays and analyzed the data as described above.

#### Electrophysiology with optogenetic inhibition of VIP-neurons

The laminar electrophysiological recordings with optogenetic inhibition of VIP-neurons (Figures 6L-N) were carried out in three VIP-Cre mice (1 male, 2 female) aged between two and six months. Surgical procedures were identical to those described above. We targeted virus injections to V1. We used a dental drill to make a small craniotomy above left V1 2.7mm lateral from the midline, 0.5mm anterior of lambda and placed a pulled borosilicate capillary containing the virus (AAV1-hSyn-SIO-stGtACR2-FusionRed, titer 1*10^13^ GC/mL, a gift from Ofer Yizhar, Addgene viral prep # 105677-AAV1) vertically above the craniotomy touching the dura. We slowly lowered the pipette to a depth of 600µm from the brain surface and slowly injected a total of 90nL at different depths (30nL at 600µm, 30nL at 400µm, 30nL at 200µm) using Nanoject III programmable injector (Drummond Scientific). After the injection, we left the pipette in place for at least 8 minutes before slowly retracting it to avoid efflux of virus. We sealed the chamber with the biocompatible adhesive Kwik-Cast (WPI). After three weeks, we made a craniotomy centered on the injection site. The mice were head-fixed on a treadmill and we performed laminar recordings, RF mappings and placed figures as described above. To test the efficacy of the approach, we first inhibited VIP-neurons in a condition with only the center grating in the RF and observed that silencing decreased visually evoked activity in V1, but that it did not influence spontaneous activity levels (Figure S7). In the main experiment with figure-ground stimuli, we activated the laser on 50% of the trials with a constant power of 5mW directed at the V1 recording site through an optical fiber (200µm thickness, FC/PC to 1.25mm Ceramic Ferrule, Thorlabs, Newton). The laser was turned on 250ms after onset of the visual stimulus and was turned off 100ms after stimulus offset. We tested the influence of optogenetic inhibition of VIP-neurons on FGM in V1 using the same statistics as described in the section on the optogenetic inhibition of higher visual areas, above.

#### Histology

To examine virus expression, we deeply anesthetized the mice with Nembutal and transcardially perfused them with phosphate buffered saline (PBS) followed by 4% paraformaldehyde (PFA) in PBS. We extracted the brain and post-fixated it overnight in 4% PFA before moving it to a PBS solution. We cut the brains into 50µm thick coronal slices and mounted them on glass slides. We imaged the slices on a Zeiss Axioplan 2 microscope (5x objective, Zeiss plan-apochromat, 0.16NA) using custom written Image-Pro Plus software and aligned the images to the Paxinos and Franklin adult mouse brain atlas (Paxinos and Franklin, 2004). To determine the location of virus expression relative to the position of cortical visual areas, we imaged the intact ex-vivo brains of M27-M31 with a wide-field microscope, averaging across 100 fluorescent images with an RFP filter using ThorCam (Thorlabs) software. To improve the signal to noise ratio, we normalized the contrast of individual images before averaging 100 images, corrected for unequal illumination by subtracting the blue channel from the red channel (FusionRed fluorescence) and smoothed the result with a 3×3 median filter. We used the population RF mapping data to determine the border of area V1, and used a bright field image of the same field of view to visualize the blood-vessel pattern and co-registered the loci of virus expression to the V1 border based on this blood vessel pattern (Figure 5B).

#### Electrophysiology during task performance

The laminar electrophysiological recordings during task performance (Figure S2) were carried out in four male C57BL/6 mice aged between two and twelve months. We implanted a head bar on the skull as described in the general methods section. Once the mice could perform the task, we performed a second surgery to chronically implant a custom-made two-shank silicone probe (Neuronexus) with 16 recording sites per shank and a spacing of 65µm between recording sites. The shanks covered 975µm, spanning the depth of V1. We made a small craniotomy above the left V1 (3mm lateral and 0.6mm anterior of lambda) and inserted the electrodes perpendicularly to the cortical surface. We lowered the electrode 1mm into the brain and allowed the brain to recover from dimpling for 15 minutes. We fixed electrode in place using dental cement and a placed a stainless-steel screw over the cerebellum to serve as a recording ground. We resumed training and recording after at least two days of recovery. We detected spikes by thresholding the signal at 3 times the standard deviation and convolved the detected spikes with a Gaussian with a standard deviation of 3.9ms (and an integral of 1). We determined the laminar position of each recording site using CSD analysis (Mitzdorf, 1985) and excluded recording sites more than 300 um above or more than 400 um below the L4/L5 border. We also excluded recording sites with a SNR below 1.5, leaving a total of 39 recording sites in the four mice for analysis. We subtracted the average baseline response from 200ms before stimulus presentation and normalized to the peak of the smoothed (moving average, 14ms span) ground condition. To measure the eye-position, we tracked the pupil of one eye contralateral to the V1 recording site using an ISCAN system and sampled it at 120Hz. We calculated the Euclidian distance from the median eye position during every trial, computed z-scores based on the distribution of distances across samples and removed all trials with absolute z-scores larger than 1.5 (this removed approximately 10% of all trials). We first measured the receptive field of the neurons at each recording site by presenting 10 by 10 deg. white (40 cd/m^2^) on black (0.02 cd/m^2^) squares in a field ranging from approximately -25 to 65 deg. horizontally and -15 to 55 deg. vertically relative to the mouse’s nose (positive values indicate the hemifield contralateral to the electrode). Squares were presented a total of 30 times in a random order. The stimulus duration was 250ms with an interval of 250ms during which the screen remained dark. We centered a 45°/50° figure on the RF and shifted it horizontally by 55° in the background condition. We included sessions for which the mouse reached a minimal accuracy of 65%, resulting in an average of 5.3 sessions and an average total of 674 trials per mouse. Due to a programming error, the figure presentation was delayed by one frame (17ms) relative to the onset of the background in one of the mice. Removing data from this mouse did not change the results.

#### Determination of the reaction times

We estimated the minimal reaction time (mRT) for all mice with sufficient trial numbers (>1000 trials) and compared mRT for contrast-orientation- and phase-defined figures. We measured the timing of the first lick in all trials (without optogenetic manipulation). Because the mice made early random licks on a substantial fraction of trials, we estimated the time point at which there were more correct than erroneous licks. We used the first of seven significantly different 10 ms bins (Chi-squared test) to estimate the mRT (Figure S3A). It was 245ms for contrast-defined figures, 255ms for orientation defined figure-ground stimuli and 315ms for phase-defined stimuli. We also determined the mRT for each mouse using the same analysis and estimated the variability (s.e.m.) by bootstrapping trials for each mouse and recalculating the mRT 1000 times. We restricted this analysis to trials with first licks occurring before 500ms after stimulus onset, as the majority of first licks occurred before this time. We determined whether mRTs in the orientation and phase-defined figure-ground task were significantly different using a paired t-test. Lick data between 0ms and 200ms is missing due to a technical reason: the Arduino used for lick detection communicated with the stimulus computer via the serial port and did not detect licks during this phase (which coincided with the grace period).

#### Determination of the efficacy of cortical silencing with hGtACR2

We performed electrophysiological recordings in two C57BL/6 mice under anesthesia to determine the efficacy of optogenetic silencing (Figures S3G,H). We injected AAV5-CaMKII-hGtACR2-eYFP in V1 of these mice. We induced anesthesia and maintained it with an intraperitoneal (i.p.) injection of urethane (1.2g/kg body weight) and chlorprothixene (8 mg/kg body weight, i.p.). Additionally, we subcutaneously (s.c.) injected atropine sulfate (0.1 mg/kg body weight) to reduce mucous secretions and dexamethasone (4mg/kg, s.c.) to prevent cortical edema. We regularly checked the depth of anesthesia with paw reflexes and gave an additional dose of urethane (200mg/kg body weight) when a response to a toe-pinch was observed. The mouse was head fixed in a stereotactic frame and the body temperature was maintained at 36.5 °C by a feedback-controlled heating pad. We applied a thin layer of adhesive to the bone, which clarifies the skull (clear skull cap). After the adhesive dried, we inserted a reference wire (Ag/Cl) between the skull and dura of the frontal cortex. We performed a small craniotomy over V1 at 0.5mm anterior of lambda and 2.7mm lateral from the midline. We inserted a linear-array recording electrode (A1×32-5mm-25-177, NeuroNexus, 32 channel probe, 25 micron spacing) in V1 and lowered it to around 1mm below the brain surface and adjusted the depth of the electrode with reference to the current source density profile to ensure coverage of all layers (Self et al., 2014). We presented a full-screen checkerboard stimulus, which was inverted after 250ms, and turned off 250ms later. On 50% of trials we inhibited neuronal activity using a DPSS Laser (BL473T3-100FC, Shanghai Lasers & Optics Century Co.) emitting blue light (wavelength 473nm) with a constant power of 5mW as a light source and directed the light at distances to the V1 electrode ranging from 0mm to 2.5mm with an optical fiber (200µm thickness). We also included trials in which the laser illuminated the brain without visual stimulation, and used these trials to correct for the light-induced artifact, subtracting neuronal activity elicited in the laser-only condition from the trials with visual stimulation plus laser stimulation. We only included recording sites with a good signal-to-noise ratio (peak visual response larger than four times the standard deviation of the baseline period).

#### Population receptive field (pRF) mapping

To localize the borders between the visual areas for the analysis of the wide-field data and to determine the placement of viral injections and electrode penetrations, we used a pRF mapping technique that was originally developed for human functional resonance magnetic imaging (Dumoulin and Wandell, 2008) (Figure S4). The mapping stimulus was constructed from a static checkerboard pattern composed of black (0cd.m^-2^) and white (40cd.m^-2^) checks of 5°x5°. The stimuli were created by presenting the checkerboard pattern within a bar-shaped aperture. The bar was 20° in width, had an orientation of 0°, 45°, 90° or 135° and was presented at different positions tiling the entire screen (Figure S4A). We used the correction for a flat screen (general methods section) and restricted the stimulus to eccentricities smaller than 70 deg. The bar stimuli were presented for 500ms, followed by an inter-stimulus interval with a grey screen for 3.6s. Each bar stimulus was presented 15 times. The calcium signal was pre-processed as described above. We took the baseline fluorescence (*F*_*0*_) as the mean fluorescence between -0.25 and 0s and the stimulus response (*F*_*n*_) as the mean fluorescence between 0.15 and 0.4s.

The mean responses of cortical pixels to the bar stimuli were fit using a forward model in which its pRF is assumed to be a 2D Gaussian envelope of the form.

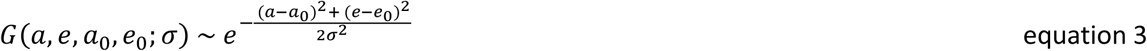

The three free parameters were the center of the Gaussian in the azimuth (a_0_) and elevation (e_0_) directions and the standard deviation (σ), which was converted to full-width at half-maximum (FWHM) by multiplying by 2.35. Gaussians were constructed with center co-ordinates ranging from -90 to +90 degrees of azimuth and -60 to +60 degrees of elevation, with a spacing of 2°. The FWHM ranged from 20° to 120° in steps of 2°. Gaussian fits with centers outside the stimulated region of space were removed and 376,431 remained. To make a predicted set of responses, we multiplied each of the Gaussian fits on a point-by-point basis with a model of each bar-stimulus. We assumed that the dark- and light-checks of the bar stimulus contribute equally to the calcium response and we used the full aperture of the stimulus in the prediction, i.e. the strength *S (a,e,i)* of stimulus *i* was 1 within the aperture and 0 outside. The predicted response (*R*) of a cortical pixel to stimulus *i* is given by:

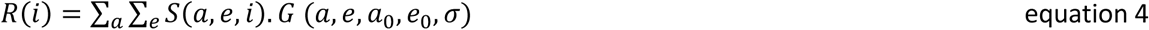

Hence, we summed across all stimulus pixels and the predicted response is proportional to the observed response *y (i)* of each cortical pixel via a single gain parameter β. We estimated β with a linear regression:

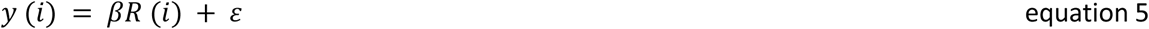

The goodness-of-fit of each predicted response was assessed using the sum-of-squares error between the observed and predicted response. The Gaussian minimizing this error term was taken as the pRF for each pixel. To reduce calculation time, we estimated the best fitting Gaussian for every other cortical pixel and linearly interpolated the results for the remaining pixels.

The resulting azimuth and elevation maps were converted into visual field sign maps using the techniques described in Garrett et al. (Garrett et al., 2014) and Sereno et al. (Sereno et al., 1995). These maps indicate whether the visual field is represented in a mirror-inverted or non-inverted fashion in cortex and we used them to determine the location of the different visual areas and the borders between them (Figures 5B, S4D).

## Supplementary Material

**Figure S1.**
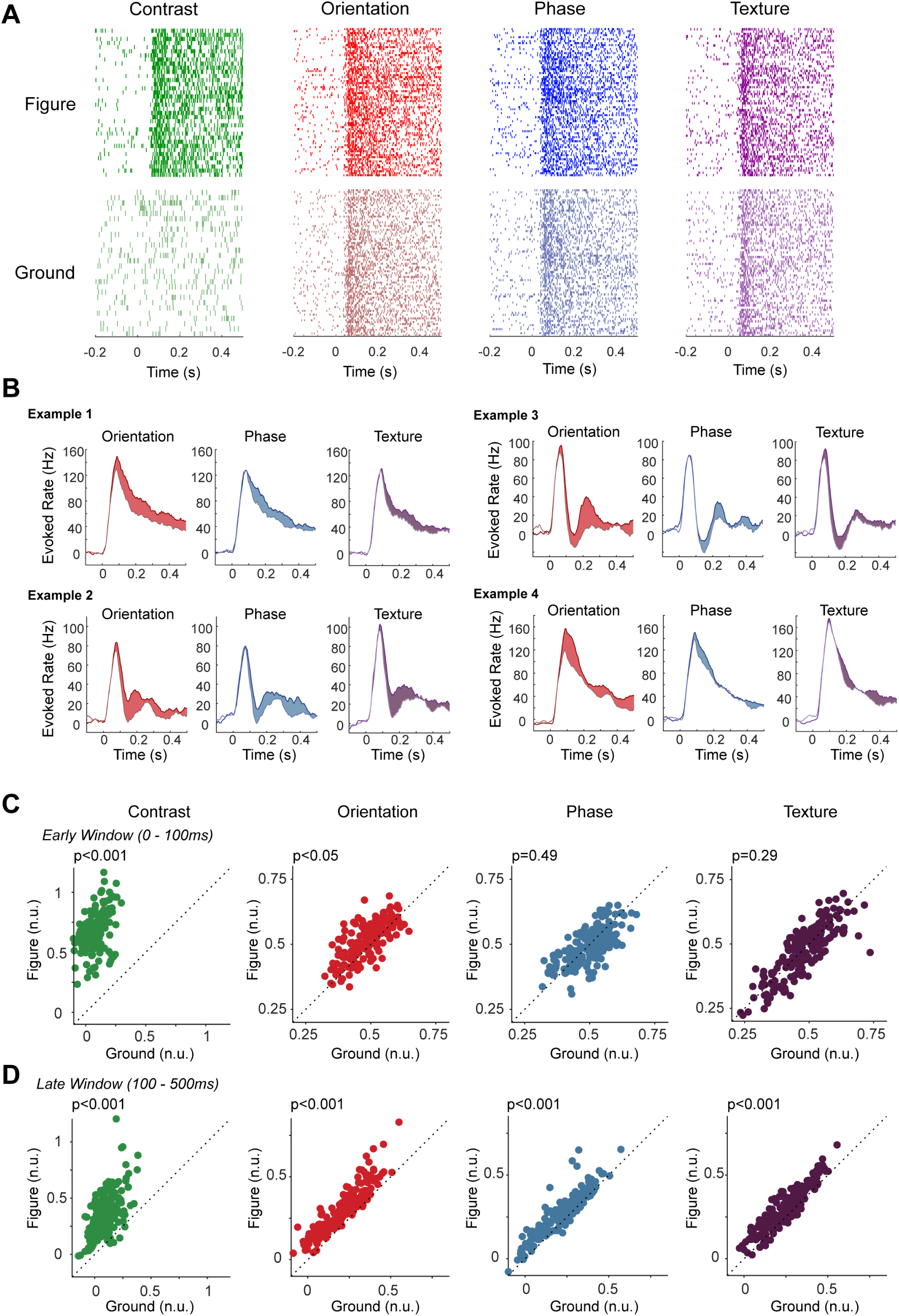
Example recording sites and variability in electrophysiological experiments, Related to Figure 2. (A) Activity elicited by the four stimuli (see Figure 1A) at an example recording site (Example 1 in panel B), shown as raster plots. (B) Activity elicited at V1 example recording sites elicited by figure-ground stimuli, where the figure was defined by a difference in orientation (left column), phase (middle column) and texture (right column). The upper traces in each plot represent the activity elicited by the figure and lower traces activity elicited by the background. The colored area in each of the panels represents the FGM. (C) Activity elicited by figure (ordinate) and background (grey screen for the contrast stimulus; abscissa) in an early (0-100ms) time window. The p-values indicate the main-effect of figure-ground as assessed by a linear mixed-effects model. (D) Same as C but in a later time window (100-500ms).

**Figure S2.**
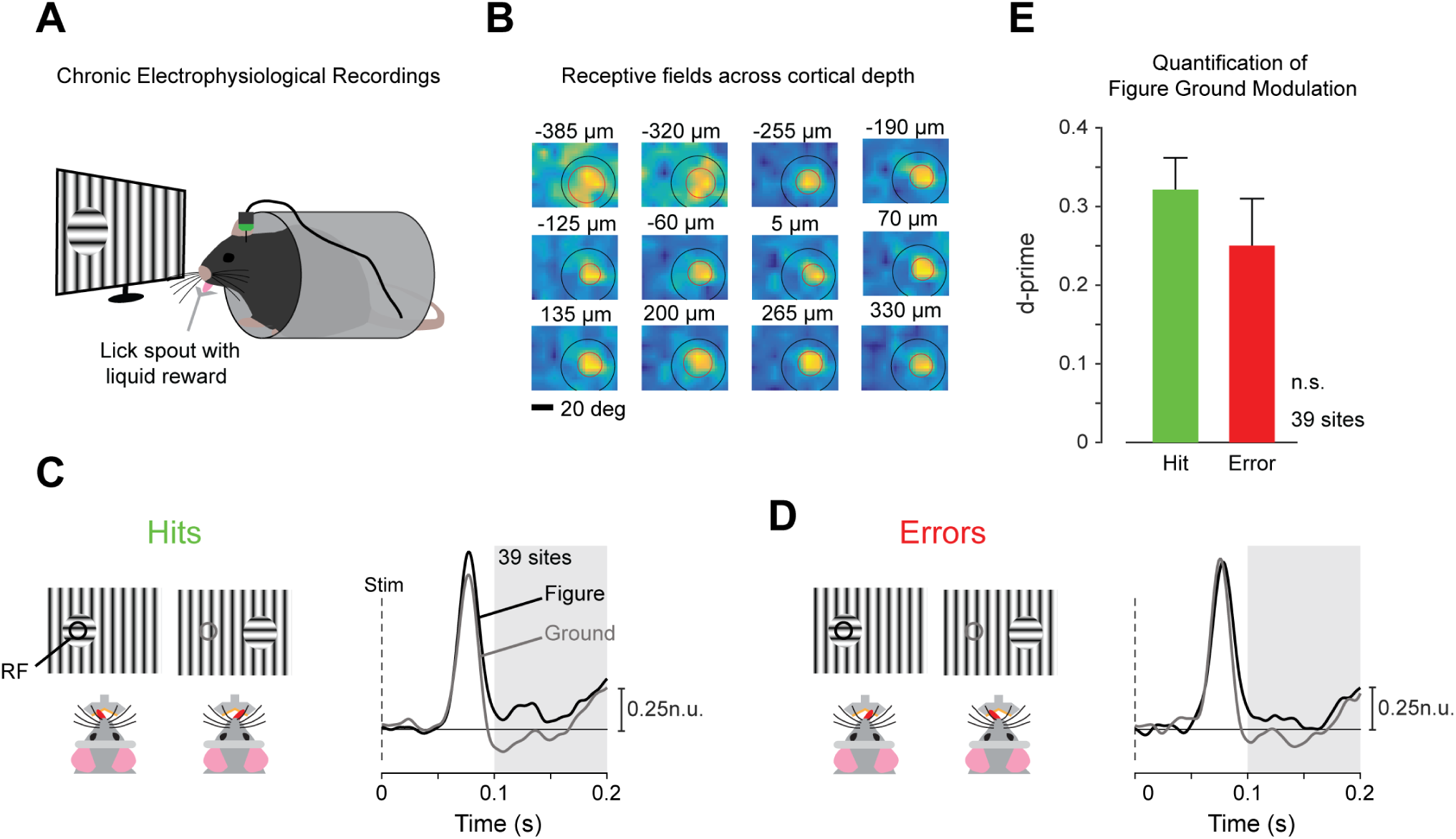
Figure-ground modulation during behavior, Related to Figure 2. (A) We recorded neural activity using implanted laminar electrodes in four mice while they indicated the side of an orientation-defined figure by licking one of two lick spouts. (B) The RFs of V1 neurons of a penetration (red circle) were aligned and fell in the center of a figure (black circle). The cortical depth relative to the layer 4/5 boundary was assessed with CSD. Negative numbers refer to cortical sites below the layer 4/5 boundary. (C) MUA response in normalized units (n.u.) averaged across 39 recording sites in V1 elicited by the figure (black) and background (grey) during trials in which the mice responded correctly. The grey area indicates the time-window used for the computation of the d-prime. (D) MUA response averaged across 39 recording sites in V1 elicited by the figure (black) and background (grey) during error trials. (E) Average FGM across 39 recording sites measured with the d-prime. Error bars, s.e.m. FGM was similar on correct and erroneous trials (paired t-test, t (38)=1.2, p=0.23, time window 100-200ms after stimulus onset).

**Figure S3.**
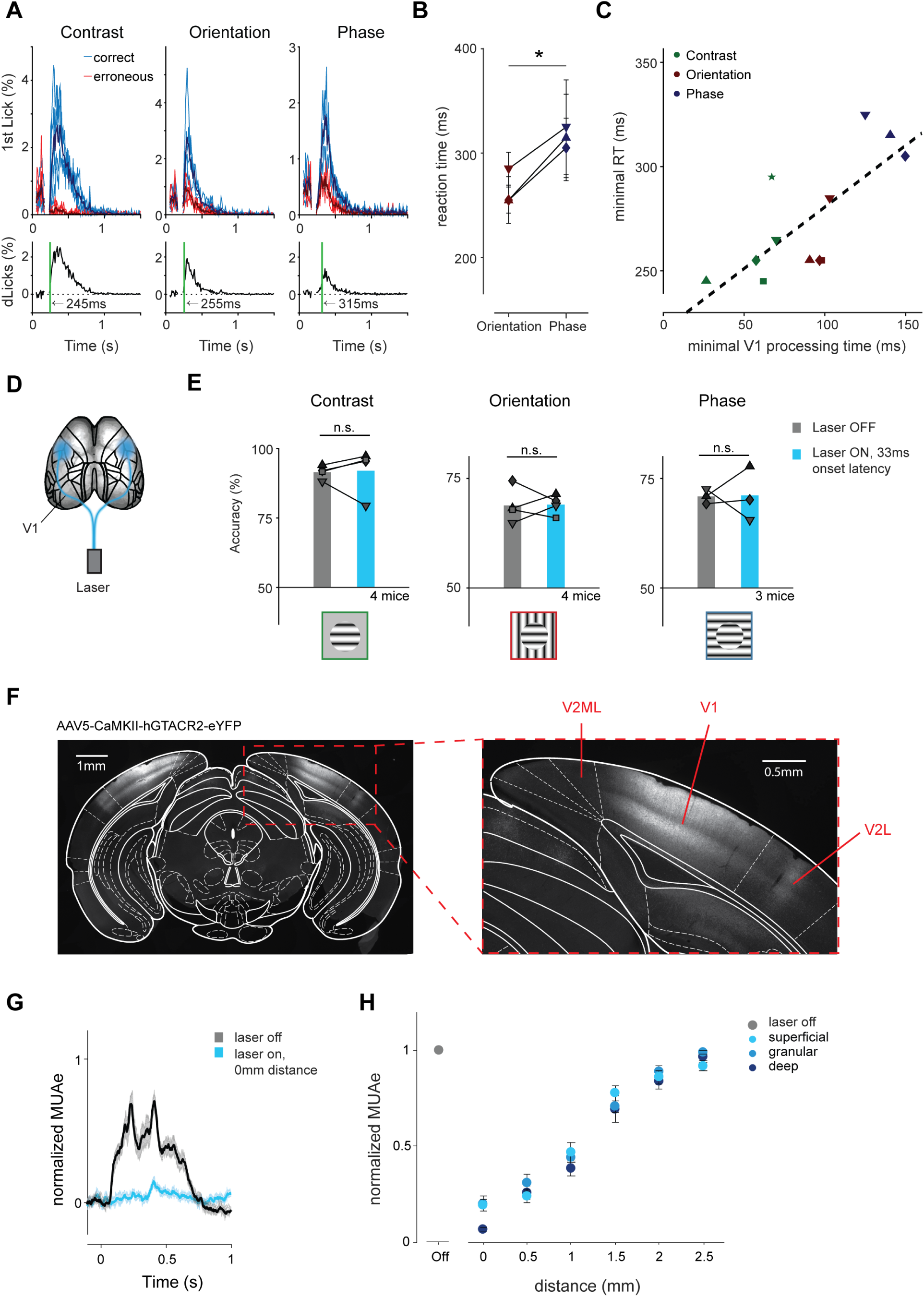
Reaction times, and reliability, specificity and spatial extent of GtACR2-mediated neuronal inhibition, Related to Figure 2. (A) upper, We determined the timing of the first lick in each trial (10ms bins) and plotted the lick count divided by the total number of trials for each cue type for correct (blue) and erroneous (red) first licks. Thin lines represent the lick distribution of individual mice and thick lines the average across mice. Lower, difference in the average distributions between correct and erroneous first licks. The green vertical line marks the minimal reaction time (mRT) for each cue. We determined mRT by taking the first of seven consecutive 10ms bins in which the correct and erroneous lick rate was significantly different (p<0.05; Chi-squared test). Lick data between 0ms and 200ms is missing due to a technical reason: the Arduino used for lick detection communicated with the stimulus computer via the serial port and did not detect licks during this phase (grace period). Time zero marks the onset of the visual stimulus. (B) The difference in mRT between orientation- and phase-defined figure-ground stimuli in the three mice that carried out both tasks was significant. *, p<0.05, t (2)=8.7, paired t-test. Error bars, s.e.m. determined by bootstrapping. (C) Correlation across mice between mRT (y-axis) and the optogenetically determined minimally necessary V1 processing time (x-axis; same data and symbols as in Figures 2D,E). The correlation coefficient was 0.75 (p<0.01). (D) To confirm that the effects of the blue laser light are caused by the shutdown of neural activity in the visual cortex, we directed the light to the somatosensory cortex of both hemispheres instead of to V1. (E) The accuracy in the figure-detection task (contrast, left), and orientation-defined (middle) and phase-defined (right) figure-ground tasks was not impaired by shining blue light on an area outside V1 in four mice (results of different mice are shown as distinct symbols; ps > 0.05 in all conditions, binomial test). (F) Example coronal slice aligned to the Paxinos and Franklin adult mouse brain atlas, showing expression pattern in mouse M49, which was injected with AAV5-CaMKII-hGtACR2-eYPF in V1 in both hemispheres. Note the strong expression within V1. The weak labelling visible in V2L is presumably caused by V1 fibers projecting to this area, which was not targeted by the laser. (G) We used laminar silicon probes to record the neuronal activity in V1 in two anesthetized mice, which had been injected with the inhibitory opsin GtACR2 in V1. Average, normalized MUA response elicited by a full-screen checkerboard stimulus in the absence (grey) and presence (blue) of optogenetic silencing with blue laser light (10mW), targeted directly at the recording site. The shaded area marks the SEM across 30 recording sites. (H) Average normalized MUA recorded at various distances from the center of the laser light and at 11 deep, 6 granular and 13 superficial recording sites. Error bars, s.e.m.

**Figure S4.**
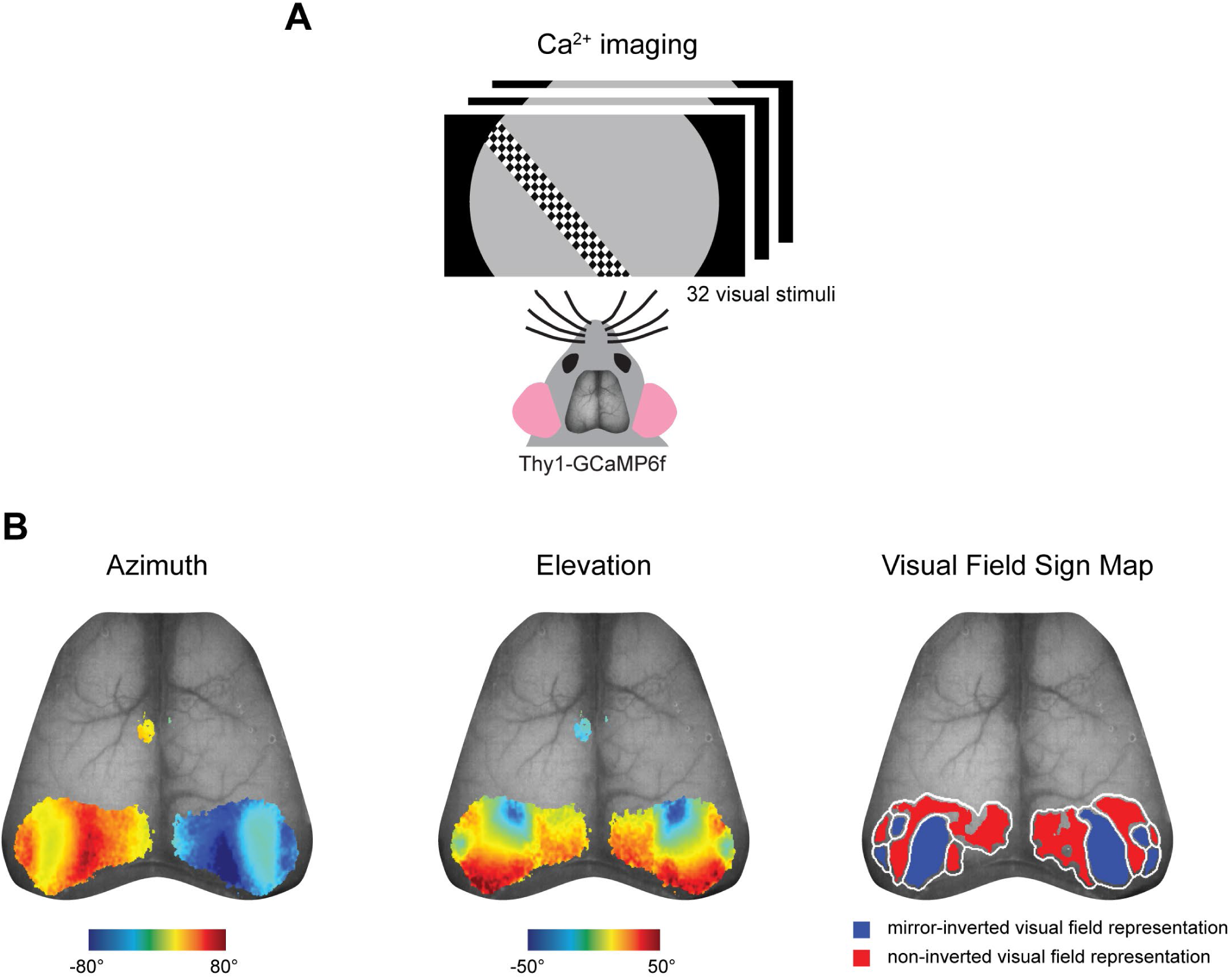
Identifying borders between cortical areas using population RF mapping, Related to Figure 3 and 5. (A) Calcium signals were imaged through the cleared skull of Thy1-GCaMP6f mice viewing stationary checkerboard bars of 32 different orientations and positions. (B) Example cortical maps showing the azimuth (left) and elevation (middle) of the best-fitting Gaussians superimposed on a wide-field image of the cortex. The maps are thresholded at a correlation value of 0.75. right, visual field sign map (non-mirror-image versus mirror-image visual field representation) generated from the azimuth and elevation maps. We used this map to determine the location and borders of visual cortical areas.

**Figure S5.**
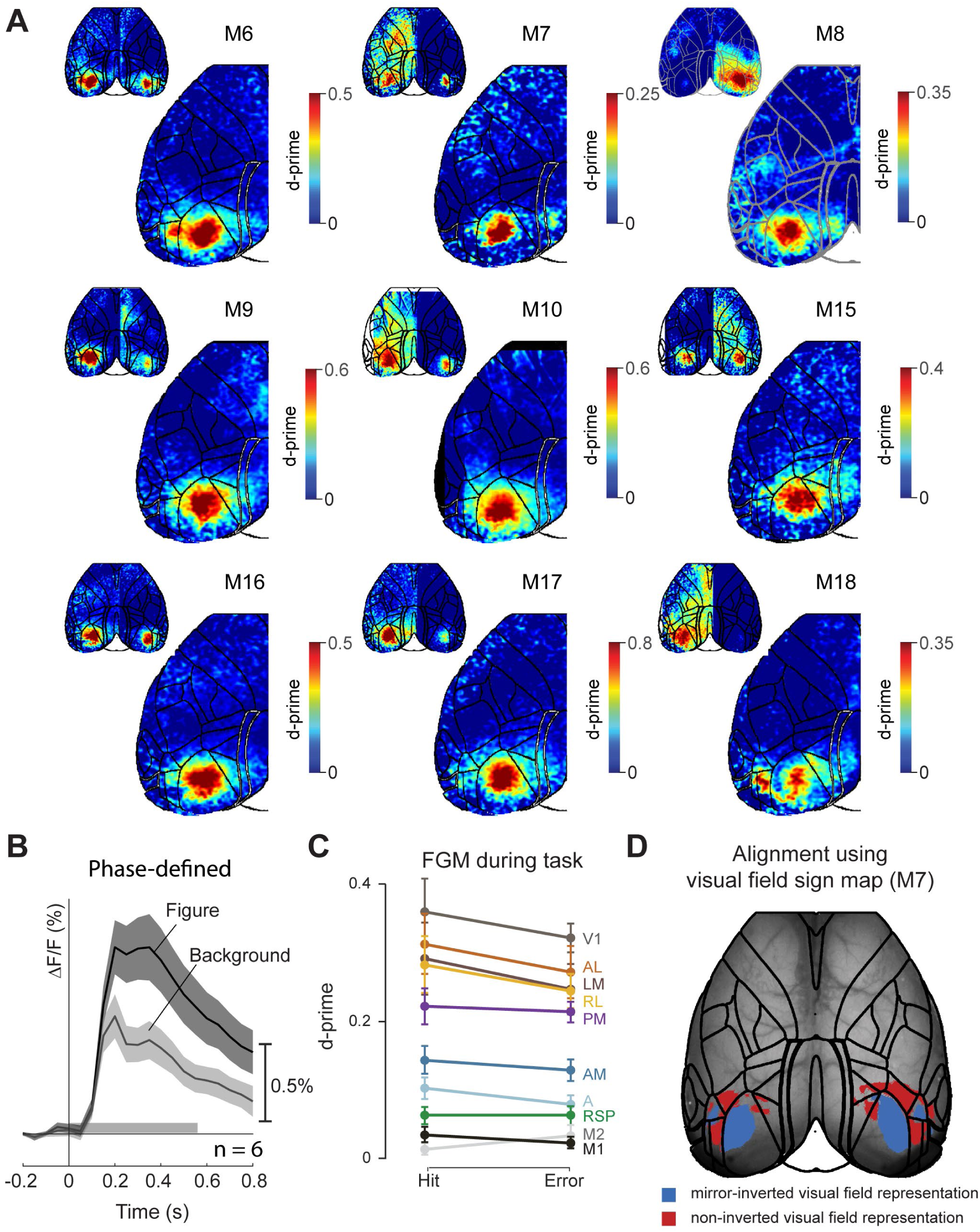
Widefield imaging during passive viewing and figure-detection, Related to Figure 3. (A) d-prime for orientation-defined figures for individual mice during passive viewing (time window 150-300ms after stimulus onset). Red regions indicate higher activity for figures than backgrounds in the contralateral hemifield. We pooled data across the two hemispheres in each mouse. Insets in the upper left show data of individual hemispheres. (B) Average V1 response elicited by a phase-defined figure (n=6 hemispheres). Black, response of cortical pixels with RFs that fell within the phase-defined figure region (black circle in the left panel of Figure 3B). Grey, response elicited by the background stimulus in same pixels. Shaded area represents s.e.m. (C) Average d-prime in the different visual areas of five mice during hits and errors. Error bars denote s.e.m. (D) Example field-sign map based on the measurement of population receptive fields in M7. Cortical areas shown in red represent the visual field in a non-inverted manner, and blue areas in a mirror-inverted fashion. We adjusted the Allen brain common coordinate framework to fit this field-sign map of individual mice by adjusting the x, y positions and scaling.

**Figure S6.**
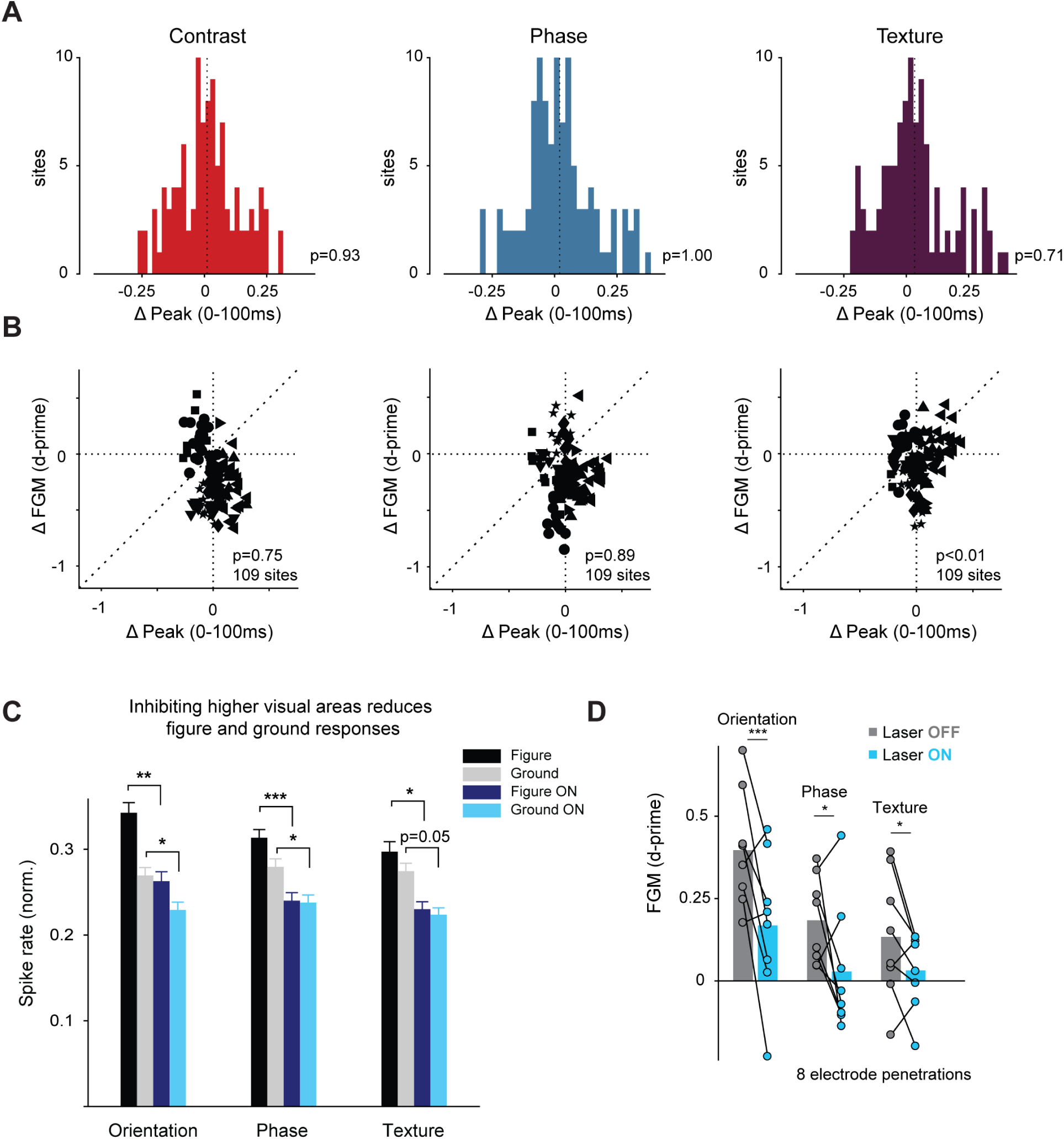
Effects of inhibiting higher visual areas on the early and late V1 response, Related to Figure 5. (A) Histogram of differences in early V1 activity (time window 0-100ms) induced by inhibiting HVAs with blue laser light. We subtracted the normalized activity in the control condition from that with the laser on. There was no significant net change in the initial V1 response for orientation-defined (left), phase-defined (middle) and textured figure-ground stimuli (right) (all ps > 0.05, paired t-test). (B) We examined whether increases or decreases in the peak response caused by inhibition of activity in HVAs predicted the effect on FGM by measuring the correlation. X-axis, change in peak response. Y-axis, change in FGM d-prime. The correlations for orientation-defined (left), phase-defined (middle), and textured figure-ground stimuli (right) were not significant (linear mixed-effects model, all ps>0.05). Hence, changes in the peak response caused by optogenetic inhibition do not predict the influence on FGM. Distinct symbols mark recording sites of different electrode penetrations. (C) Average activity (time window 100-500ms after stimulus presentation) of 109 sites normalized to the peak response. Inhibition of HVAs reduced the responses to figures (compare black and dark blue) and backgrounds (compare grey and light blue) for all figure-ground stimuli (*, p<0.05; **, p<0.01; ***, p<0.001, linear mixed-effects model, see Methods). Error bars, s.e.m. (D) Average d-prime values of FGM per recording session without (grey) and with optogenetic inhibition (blue) of HVAs for different figure-ground stimuli. We made 8 electrode penetrations in 5 mice. Bars show the average d-prime across sessions.

**Figure S7.**
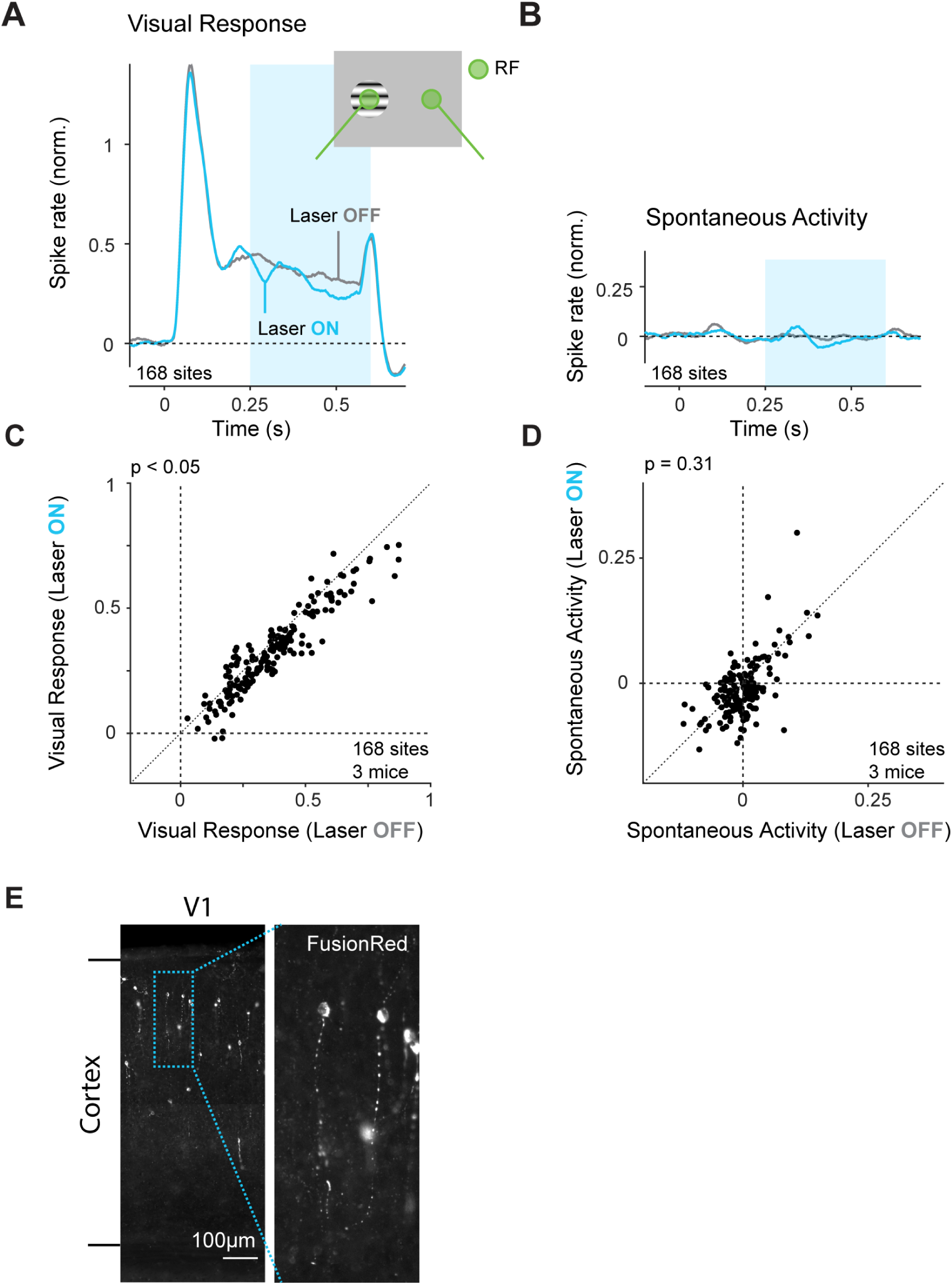
Inhibiting VIP-neurons decreases the visual response of V1 neurons, Related to Figure 6. (A,B) We presented circular figures on a grey background, where the figure was either centered on the RF of the recorded V1 neurons (A) or placed 50-60 degrees away (B). We optogenetically inhibited VIP-neurons in 50% of the trials using blue laser light, starting 250ms after stimulus onset. Average MUA across 168 recording sites (8 penetrations, 3 mice) with (blue) and without (grey) inhibition of VIP-neurons. (C,D) Inhibition of VIP-neurons significantly reduced the visually evoked activity in V1 (C) (p<0.05, linear mixed-effects model, time window 250-500ms), but not spontaneous activity (D) (p=0.31, linear mixed-effects model). (E) left, Image of coronal section of V1, spanning from the bottom to the top of the cortex. Expression of optogenetic construct is visualized by the reporter molecule FusionRed. right, Enlarged view of the blue rectangle.

**Table S1.** Accuracies of mice in the different tasks

| Mouse | Genotype | Experiment | Contrast |  | Orientation |  | Phase |  | Texture |  |
| --- | --- | --- | --- | --- | --- | --- | --- | --- | --- | --- |
|  |  |  | Accu-<br>racy<sup>b)</sup> | Number of<br>Trials<sup>c)</sup> | Accu-<br>racy<sup>b)</sup> | Number of<br>Trials<sup>c)</sup> | Accu-<br>racy<sup>b)</sup> | Number of<br>Trials<sup>c)</sup> | Accu-<br>racy<sup>b)</sup> | Number of<br>Trials<sup>c)</sup> |
| M1<br>(male) | C57BL/6<sup>a)</sup> | Optogenetics<br>(Fig. 1C, 2C-E) | 94.62%* | 966 (1012),<br>384 (399) | 71.98%* | 1035 (1306),<br>399 (430) | 70.49%* | 1484 (1890),<br>450 (498) | 66.84%* | 1942<br>(2362) |
|  |  | Optogenetics ctrl.<br>(Fig. S3B) | 92.04%* | 226 (234),<br>94 (94) | 74.47%* | 427 (475),<br>146 (152) | 69.23%* | 247 (315),<br>67 (81) | - | - |
| M2<br>(male) | C57BL/6<sup>a)</sup> | Optogenetics<br>(Fig. 1C, 2C-E) | 91.54%* | 1028 (1063),<br>315 (344) | 66.84%* | 1565 (1745),<br>452 (489) | - | - | - | - |
|  |  | Optogenetics ctrl.<br>(Fig. S3B) | 91.67%* | 72 (73),<br>44 (44) | 67.89%* | 246 (273),<br>50 (62) | - | - | - | - |
| M3<br>(male) | C57BL/6<sup>a)</sup> | Optogenetics<br>(Fig. 1C, 2C-E) | 96.57%* | 1108 (1190),<br>657 (725) | 71.44%* | 1348 (1602),<br>439 (496) | 66.42%* | 1635 (2378),<br>439 (539) | 64.20%* | 1461<br>(2891) |
|  |  | Optogenetics ctrl.<br>(Fig. S3B) | 94.01%* | 117 (119),<br>71 (76) | 68.21%* | 409 (480),<br>84 (104) | 70.96%* | 124 (183),<br>18 (28) | - | - |
| M4<br>(female) | C57BL/6<sup>a)</sup> | Optogenetics<br>(Fig. 1C, 2C-E) | 80.07%* | 1385 (1773),<br>412 (689) | - | - | - | - | - | - |
| M5<br>(male) | C57BL/6<sup>a)</sup> | Optogenetics<br>(Fig. 1C, 2C-E) | 87.34%* | 1777 (1861),<br>527 (732) | 69.41%* | 2105 (2866),<br>363 (562) | 68.21%* | 1809 (2776),<br>395 (754) | - | - |
|  |  | Optogenetics ctrl.<br>(Fig. S3B) | 88.12%* | 202 (211),<br>131 (134) | 64.83%* | 290 (390),<br>202 (274) | 72.57%* | 113 (190),<br>29 (49) | - | - |
| M6<br>(male) | Thy1-GCaMP6f | Wide-field imaging<br>(Fig. 1C, 3, S5) | 90.57%* | 53 (119) | 84.35%* | 147 (325) | 70.66%* | 167 (296) | - | - |
| M7<br>(male) | Thy1-GCaMP6f | Wide-field imaging<br>(Fig. 1C, 3, S4, S5) | 91.81%* | 220 (273) | 71.30%* | 553 (720) | - | - | - | - |
| M8<br>(male) | Thy1-GCaMP6f | Wide-field imaging<br>(Fig. 1C, 3, S5) | 93.28%* | 134 (160) | 65.71%* | 382 (470) | - | - | - | - |
| M9<br>(female) | Thy1-GCaMP6f | Wide-field imaging<br>(Fig. 1C, 3, S5) | 75.82%* | 91 (109) | 73.54%* | 257 (325) | 72.39%* | 239 (295) | - | - |
| M10<br>(male) | Thy1-GCaMP6f | Wide-field imaging<br>(Fig. 1C, 3, S5) | 95.14%* | 144 (179) | 78.47%* | 288 (388) | 73.75%* | 339 (431) | 62.50%* | 664 (837) |
| M11<br>(male) | C57BL/6 | Electrophysiology<br>(Fig. 1C, S2) | - | - | 64.63%* | 759 (912) | - | - | - | - |
| M12<br>(male) | C57BL/6 | Electrophysiology<br>(Fig. 1C, S2) | - | - | 75.64%* | 420 (711) | - | - | - | - |
| M13<br>(male) | C57BL/6 | Electrophysiology<br>(Fig. 1C, S2) | - | - | 69.85%* | 537 (625) | - | - | - | - |
| M14<br>(male) | C57BL/6 | Electrophysiology<br>(Fig. 1C, S2) | - | - | 78.73%* | 442 (1202) | - | - | - | - |

**Table S2.**
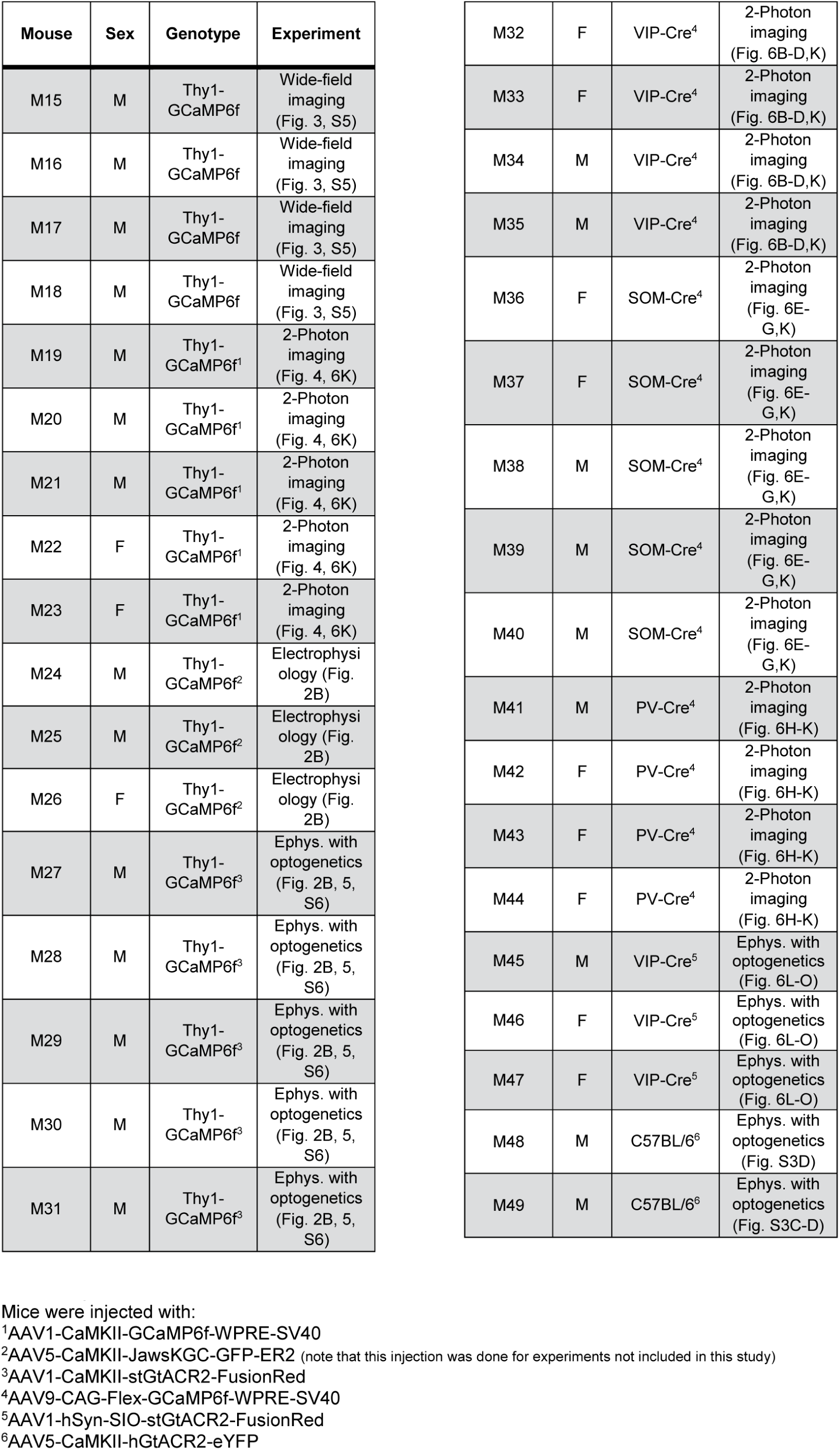
Overview of mice used in passive experiments

## References

Adesnik, H., Bruns, W., Taniguchi, H., Huang, Z.J., and Scanziani, M. (2012). A neural circuit for spatial summation in visual cortex. Nature 490, 226–231.

Allen, W.E., Kauvar, I. V., Chen, M.Z., Richman, E.B., Yang, S.J., Chan, K., Gradinaru, V., Deverman, B.E., Luo, L., and Deisseroth, K. (2017). Global Representations of Goal-Directed Behavior in Distinct Cell Types of Mouse Neocortex. Neuron 94, 891–907.

Atallah, B. V., Bruns, W., Carandini, M., and Scanziani, M. (2012). Parvalbumin-Expressing Interneurons Linearly Transform Cortical Responses to Visual Stimuli. Neuron 73, 159–170.

Baldauf, D., and Deubel, H. (2010). Attentional landscapes in reaching and grasping. Vision Res. 50, 999–1013.

Bhatt, R., Carpenter, G.A., and Grossberg, S. (2007). Texture segregation by visual cortex: Perceptual grouping, attention, and learning. Vision Res. 47, 3173–3211.

Chelazzi, L., Miller, E.K., Duncan, J., and Desimone, R. (1993). A neural basis for visual search in inferior temporal cortex. Nature 363, 345–347.

Chen, M., Yan, Y., Gong, X., Gilbert, C.D., Liang, H., and Li, W. (2014). Incremental Integration of Global Contours through Interplay between Visual Cortical Areas. Neuron 82, 682–694.

Cowey, A., and Stoerig, P. (1995). Blindsight in monkeys. Nature 373, 247–249.

Cowey, A., and Weiskrantz, L. (1963). A Perimetric Study of Visual Field Defects in Monkeys. Q. J. Exp. Psychol. 15, 91–115.

D’Souza, R.D., Meier, A.M., Bista, P., Wang, Q., and Burkhalter, A. (2016). Recruitment of inhibition and excitation across mouse visual cortex depends on the hierarchy of interconnecting areas. Elife DOI: 10.7554/eLife.19332.

Dana, H., Chen, T.-W., Hu, A., Shields, B.C., Guo, C., Looger, L.L., Kim, D.S., and Svoboda, K. (2014). Thy1-GCaMP6 Transgenic Mice for Neuronal Population Imaging In Vivo. PLoS One DOI: 10.1371/journal.pone.0108697.

Desimone, R., and Duncan, J. (1995). Neural mechanisms of selective visual attention. Annu. Rev. Neurosci. 18, 193–222.

Dipoppa, M., Ranson, A., Krumin, M., Pachitariu, M., Carandini, M., and Harris, K.D. (2018). Vision and Locomotion Shape the Interactions between Neuron Types in Mouse Visual Cortex. Neuron 98, 602–615.

Dumoulin, S.O., and Wandell, B.A. (2008). Population receptive field estimates in human visual cortex. Neuroimage 39, 647–660.

Duncan, J., Humphreys, G.K., and Ward, R. (1997). Competitive brain activity in visual attention. Curr. Opin. Neurobiol. 7, 255–261.

Erisken, S., Vaiceliunaite, A., Jurjut, O., Fiorini, M., Katzner, S., and Busse, L. (2014). Effects of locomotion extend throughout the mouse early visual system. Curr. Biol. 24, 2899–2907.

Gamanut, R., Kennedy, H., Toroczkai, Z., Ercsey-Ravasz, M., Van Essen, D.C., Knoblauch, K., and Burkhalter, A. (2018). The Mouse Cortical Connectome, Characterized by an Ultra-Dense Cortical Graph, Maintains Specificity by Distinct Connectivity Profiles. Neuron 97, 698–715.

Garcia Del Molino, L.C., Yang, G.R., Mejias, J.F., and Wang, X.J. (2017). Paradoxical response reversal of top-down modulation in cortical circuits with three interneuron types. Elife DOI: 10.7554/eLife.29742.

Garrett, M.E., Nauhaus, I., Marshel, J.H., and Callaway, E.M. (2014). Topography and Areal Organization of Mouse Visual Cortex. J. Neurosci. 34, 12587–12600.

Govorunova, E.G., Sineshchekov, O.A., Janz, R., Liu, X., and Spudich, J.L. (2015). Natural light-gated anion channels: A family of microbial rhodopsins for advanced optogenetics. Science 349, 647–650.

Guo, Z. V., Li, N., Huber, D., Ophir, E., Gutnisky, D., Ting, J.T., Feng, G., and Svoboda, K. (2014). Flow of cortical activity underlying a tactile decision in mice. Neuron 81, 179–194.

Harris, K.D., and Shepherd, G.M.G. (2015). The neocortical circuit: themes and variations. Nat. Neurosci. 18, 170–181.

Huh, C.Y.L., Peach, J.P., Bennett, C., Vega, R.M., and Hestrin, S. (2018). Feature-Specific Organization of Feedback Pathways in Mouse Visual Cortex. Curr. Biol. 28, 114–120.

Hupé, J.M., James, A.C., Payne, B.R., Lomber, S.G., Girard, P., and Bullier, J. (1998). Cortical feedback improves discrimination between figure and background by V1, V2 and V3 neurons. Nature 394, 784–787.

Ji, X. -y., Zingg, B., Mesik, L., Xiao, Z., Zhang, L.I., and Tao, H.W. (2015). Thalamocortical innervation pattern in mouse auditory and visual cortex: laminar and cell-type specificity. Cereb. Cortex 26, 2612–2625.

Juavinett, A.L., and Callaway, E.M. (2015). Pattern and Component Motion Responses in Mouse Visual Cortical Areas. Curr. Biol. 25, 1759–1764.

Kar, K., Kubilius, J., Schmidt, K., Issa, E.B., and DiCarlo, J.J. (2019). Evidence that recurrent circuits are critical to the ventral stream’s execution of core object recognition behavior. Nat. Neurosci. 22, 974–983.

Karnani, M.M., Jackson, J., Ayzenshtat, I., Hamzehei Sichani, A., Manoocheri, K., Kim, S., and Yuste, R. (2016). Opening holes in the blanket of inhibition: localized lateral disinhibition by VIP interneurons. J. Neurosci. 36, 3471–3480.

Keller, A.J., Roth, M.M., and Scanziani, M. (2020). Feedback generates a second receptive field in neurons of the visual cortex. Nature DOI: 10.1038/s41586-020-2319-4.

van Kerkoerle, T., Self, M.W., and Roelfsema, P.R. (2017). Layer-specificity in the effects of attention and working memory on activity in primary visual cortex. Nat. Commun. DOI: 10.1038/ncomms13804.

Kirchner, H., and Thorpe, S.J. (2006). Ultra-rapid object detection with saccadic eye movements: Visual processing speed revisited. Vision Res. 46, 1762–1776.

Klink, P.C., Dagnino, B., Gariel-Mathis, M.-A.A.M.-A., and Roelfsema, P.R. (2017). Distinct Feedforward and Feedback Effects of Microstimulation in Visual Cortex Reveal Neural Mechanisms of Texture Segregation. Neuron 95, 209–220.

Lamme, V.A. (1995). The neurophysiology of figure-ground segregation in primary visual cortex. J. Neurosci. 15, 1605–1615.

Lamme, V.A.F., and Roelfsema, P.R.R. (2000). The distinct modes of vision offered by feedforward and recurrent processing. Trends Neurosci. 23, 571–579.

Li, Z. (1999). Contextual influences in V1 as a basis for pop out and asymmetry in visual search. Proc. Natl. Acad. Sci. 96, 10530–10535.

Li, F., Jiang, W., Wang, T.-Y., Xie, T., and Yao, H. (2018). Phase-specific Surround suppression in Mouse Primary Visual Cortex Correlates with Figure Detection Behavior Based on Phase Discontinuity. Neuroscience 379, 359–374.

Mahn, M., Gibor, L., Patil, P., Cohen-Kashi Malina, K., Oring, S., Printz, Y., Levy, R., Lampl, I., and Yizhar, O. (2018). High-efficiency optogenetic silencing with soma-targeted anion-conducting channelrhodopsins. Nat. Commun. DOI: 10.1038/s41467-018-06511-8.

Manita, S., Suzuki, T., Homma, C., Matsumoto, T., Odagawa, M., Yamada, K., Ota, K., Matsubara, C., Inutsuka, A., Sato, M., et al. (2015). A Top-Down Cortical Circuit for Accurate Sensory Perception. Neuron 86, 1304–1316.

Marshel, J.H., Garrett, M.E., Nauhaus, I., and Callaway, E.M. (2011). Functional specialization of seven mouse visual cortical areas. Neuron 72, 1040–1054.

Mitchell, J.F., Sundberg, K.A., and Reynolds, J.H. (2007). Differential attention-dependent response modulation across cell classes in macaque visual area V4. Neuron 55, 131–141.

Mitzdorf, U. (1985). Current source-density method and application in cat cerebral cortex: investigation of evoked potentials and EEG phenomena. Physiol. Rev. 65, 37–100.

Nassi, J.J., Lomber, S.G., and Born, R.T. (2013). Corticocortical Feedback Contributes to Surround Suppression in V1 of the Alert Primate. J. Neurosci. 33, 8504–8517.

Niell, C.M., and Stryker, M.P. (2010). Modulation of Visual Responses by Behavioral State in Mouse Visual Cortex. Neuron 65, 472–479.

O’Craven, K.M., Downing, P.E., and Kanwisher, N. (1999). fMRI evidence for objects as the units of attentional selection. Nature 401, 584–587.

Oh, S.W., Harris, J.A., Ng, L., Winslow, B., Cain, N., Mihalas, S., Wang, Q., Lau, C., Kuan, L., Henry, A.M., et al. (2014). A mesoscale connectome of the mouse brain. Nature 508, 207–214.

Osakada, F., Mori, T., Cetin, A.H., Marshel, J.H., Virgen, B., and Callaway, E.M. (2011). New Rabies Virus Variants for Monitoring and Manipulating Activity and Gene Expression in Defined Neural Circuits. Neuron 71, 617–631.

Owen, S.F., Liu, M.H., and Kreitzer, A.C. (2019). Thermal constraints on in vivo optogenetic manipulations. Nat. Neurosci. 22, 1061–1065.

Pakan, J.M.P., Dylda, E., Currie, S.P., Coutts, C.A., Rochefort, N.L., Lowe, S.C., and Keemink, S.W. (2016). Behavioral-state modulation of inhibition is context-dependent and cell type specific in mouse visual cortex. Elife DOI: 10.7554/eLife.14985.

Paxinos, G., and Franklin, K.B.J. (2004). Mouse Brain in Stereotaxic Coordinates.

Pfeffer, C.K., Xue, M., He, M., Huang, Z.J., and Scanziani, M. (2013). Inhibition of inhibition in visual cortex: the logic of connections between molecularly distinct interneurons. Nat. Neurosci. 16, 1068–1076.

Pho, G.N., Goard, M.J., Woodson, J., Crawford, B., and Sur, M. (2018). Task-dependent representations of stimulus and choice in mouse parietal cortex. Nat. Commun. DOI: 10.1038/s41467-018-05012-y.

Pi, H.J., Hangya, B., Kvitsiani, D., Sanders, J.I., Huang, Z.J., and Kepecs, A. (2013). Cortical interneurons that specialize in disinhibitory control. Nature 503, 521–524.

Pnevmatikakis, E.A., and Giovannucci, A. (2017). NoRMCorre: An online algorithm for piecewise rigid motion correction of calcium imaging data. J. Neurosci. Methods 291, 83–94.

Pnevmatikakis, E.A., Soudry, D., Gao, Y., Machado, T.A., Merel, J., Pfau, D., Reardon, T., Mu, Y., Lacefield, C., Yang, W., et al. (2016). Simultaneous Denoising, Deconvolution, and Demixing of Calcium Imaging Data. Neuron 89, 285–299.

Poort, J., Raudies, F., Wannig, A., Lamme, V.A.F., Neumann, H., and Roelfsema, P.R. (2012). The Role of Attention in Figure-Ground Segregation in Areas V1 and V4 of the Visual Cortex. Neuron 75, 143–156.

Poort, J., Self, M.W., Van Vugt, B., Malkki, H., and Roelfsema, P.R. (2016). Texture Segregation Causes Early Figure Enhancement and Later Ground Suppression in Areas V1 and V4 of Visual Cortex. Cereb. Cortex 26, 3964–3976.

Pöppel, E., Held, R., and Frost, D. (1973). Residual Visual Function after Brain Wounds involving the Central Visual Pathways in Man. Nature 243, 295–296.

Prusky, G.T., and Douglas, R.M. (2004). Characterization of mouse cortical spatial vision. Vision Res. 44, 3411–3418.

Resulaj, A., Ruediger, S., Olsen, S.R., and Scanziani, M. (2018). First spikes in visual cortex enable perceptual discrimination. Elife DOI: 10.7554/eLife.34044.001.

Roelfsema, P.R. (2006). Cortical algorithms for perceptual grouping. Annu. Rev. Neurosci. 29, 203–227.

Roelfsema, P.R., and Houtkamp, R. (2011). Incremental grouping of image elements in vision. Attention, Perception, Psychophys. 73, 2542–2572.

Roelfsema, P.R., and de Lange, F.P. (2016). Early Visual Cortex as a Multiscale Cognitive Blackboard. Annu. Rev. Vis. Sci. 2, 131–151.

Roelfsema, P.R., Lamme, V.A.F., Spekreijse, H., and Bosch, H. (2002). Figure-ground segregation in a recurrent network architecture. J. Cogn. Neurosci. 14, 525–537.

Sachidhanandam, S., Sreenivasan, V., Kyriakatos, A., Kremer, Y., and Petersen, C.C.H.H. (2013). Membrane potential correlates of sensory perception in mouse barrel cortex. Nat. Neurosci. 16, 1671–1677.

Sanders, M.D., Warrington, E.K., Marshall, J., and Wieskrantz, L. (1974). “Blindsight”: Vision in a Field Defect. Lancet 303, 707–708.

Scheffe, H. (1956). A “Mixed Model” for the Analysis of Variance. Ann. Math. Stat. 27, 23–36.

Schmid, M.C., Mrowka, S.W., Turchi, J., Saunders, R.C., Wilke, M., Peters, A.J., Ye, F.Q., and Leopold, D.A. (2010). Blindsight depends on the lateral geniculate nucleus. Nature 466, 373–377.

Schnabel, U.H., Bossens, C., Lorteije, J.A.M., Self, M.W., Op de Beeck, H., and Roelfsema, P.R. (2018). Figure-ground perception in the awake mouse and neuronal activity elicited by figure-ground stimuli in primary visual cortex. Sci. Rep. DOI: 10.1038/s41598-018-36087-8.

Scholte, H.S., Jolij, J., Fahrenfort, J.J., and Lamme, V.A.F. (2008). Feedforward and Recurrent Processing in Scene Segmentation: Electroencephalography and Functional Magnetic Resonance Imaging. J. Cogn. Neurosci. 20, 2097–2109.

Self, M.W., Lorteije, J. a. M., Vangeneugden, J., van Beest, E.H., Grigore, M.E., Levelt, C.N., Heimel, J. a., and Roelfsema, P.R. (2014). Orientation-Tuned Surround Suppression in Mouse Visual Cortex. J. Neurosci. 34, 9290–9304.

Self, M.W., Possel, J.K., Jeurissen, D., Roelfsema, P.R., Peters, J.C., Reithler, J., Goebel, R., Ris, P., Reddy, L., Claus, S., et al. (2016). The Effects of Context and Attention on Spiking Activity in Human Early Visual Cortex. PLoS Biol. DOI: 10.1371/journal.pbio.1002420.

Self, M.W., Jeurissen, D., van Ham, A.F., van Vugt, B., Poort, J., and Roelfsema, P.R. (2019). The Segmentation of Proto-Objects in the Monkey Primary Visual Cortex. Curr. Biol. 29, 1019–1029.

Sereno, M., Dale, A., Reppas, J., Kwong, K., Belliveau, J., Brady, T., Rosen, B., and Tootell, R. (1995). Borders of multiple visual areas in humans revealed by functional magnetic resonance imaging. Science 268, 889–893.

Sherman, S.M. (2016). Thalamus plays a central role in ongoing cortical functioning. Nat. Neurosci. 19, 533–541.

Smith, I.T., Townsend, L.B., Huh, R., Zhu, H., and Smith, S.L. (2017). Stream-dependent development of higher visual cortical areas. Nat. Neurosci. 20, 200–208.

Supèr, H., Spekreijse, H., and Lamme, V.A.F. (2001). Two distinct modes of sensory processing observed in monkey primary visual cortex. Nat. Neurosci. 4, 304–310.

Thorpe, S., Fize, D., and Marlot, C. (1996). Speed of processing in the human visual system. Am. J. Ophthalmol. 122, 608–609.

Tremblay, R., Lee, S., and Rudy, B. (2016). GABAergic Interneurons in the Neocortex: From Cellular Properties to Circuits. Neuron 91, 260–292.

Wall, N.R., de la Parra, M., Sorokin, J.M., Taniguchi, H., Huang, Z.J., and Callaway, E.M. (2016). Brain-wide maps of synaptic input to cortical interneurons. J. Neurosci. 36, 4000–4009.

Wang, Q., Gao, E., and Burkhalter, A. (2011). Gateways of ventral and dorsal streams in mouse visual cortex. J. Neurosci. 31, 1905–1918.

Wang, Q., Sporns, O., and Burkhalter, A. (2012). Network Analysis of Corticocortical Connections Reveals Ventral and Dorsal Processing Streams in Mouse Visual Cortex. J. Neurosci. 32, 4386–4399.

Weiskrantz, L., Warrington, E.K., Sanders, M.D., and Marshall, J. (1974). Visual capacity in the hemianopic field following a restricted occipital ablation the relationship between attention and visual experience. Brain 97, 709–728.

Williams, L.E., and Holtmaat, A. (2019). Higher-Order Thalamocortical Inputs Gate Synaptic Long-Term Potentiation via Disinhibition. Neuron 101, 91–102.

Wokke, M.E., Sligte, I.G., Steven Scholte, H., and Lamme, V.A.F. (2012). Two critical periods in early visual cortex during figure-ground segregation. Brain Behav. 2, 763–777.

Yamins, D.L.K., and DiCarlo, J.J. (2016). Using goal-driven deep learning models to understand sensory cortex. Nat. Neurosci. 19, 356–365.

Yang, W., Carrasquillo, Y., Hooks, B.M., Nerbonne, J.M., and Burkhalter, A. (2013). Distinct balance of excitation and inhibition in an interareal feedforward and feedback circuit of mouse visual cortex. J. Neurosci. 33, 17373–17384.

Zhang, S., Xu, M., Kamigaki, T., Hoang Do, J.P.P., Chang, W.-C., Jenvay, S., Miyamichi, K., Luo, L., and Dan, Y. (2014). Long-range and local circuits for top-down modulation of visual cortex processing. Science 345, 660–665.

